# Comprehensive benchmarking of somatic structural variant detection at ultra-low allele fractions

**DOI:** 10.1101/2025.09.18.677206

**Authors:** Yuwei Zhang, Adam C. English, Luis F. Paulin, Christopher M Grochowski, Surabhi Maheshwari, Taralynn Mack, Michele Berselli, Alexander D Veit, Yilei Fu, SMaHT SV working group, Peter J Park, Fritz J Sedlazeck

**Affiliations:** Department of Biomedical Informatics, Harvard Medical School, Boston, MA, USA; Human Genome Sequencing Center, Baylor College of Medicine, Houston, TX, USA; Spiral Genetics, Seattle, WA 98101, USA; Department of Genome Sciences, University of Washington School of Medicine, Seattle, WA, USA; Department of Molecular and Human Genetics, Baylor College of Medicine, TX, USA; Department of Computer Science, Rice University, 6100 Main Street, Houston, TX, USA

**Author notes:** equal contribution (Yuwei Zhang, Adam C. English).

## Abstract

Postzygotic mosaicism gives rise to somatic structural variants (SVs) at ultra-low variant allele fractions (VAFs), which pose challenges for detection due to the high-coverage sequencing required and noise introduced by sequencing artifacts. Although somatic SV detection has been extensively studied in cancer, these studies are not directly applicable to the study of tissue mosaicism, as they rely on matched normals, target higher VAF ranges, and are enriched for different types of SVs.

We present comprehensive benchmark data and best practices for non-cancer somatic SV detection. We created a synthetic mosaic sample by combining six HapMap individuals at varying proportions, generating allele fractions as low as 0.25%. This sample was sequenced to ∼2,300x total coverage using Illumina, PacBio, and Nanopore technologies across multiple sequencing centers. A high-confidence benchmark SV set containing over 21,000 pseudo-somatic insertions and deletions ≥50bp was derived from haplotype-resolved assemblies.

We evaluated 12 SV discovery pipelines and identified caller-specific strengths and sequencing platform-specific shortcomings. We find that short read-based approaches show reduced recall for insertions and repeat-associated SVs, whereas long-read sequencing achieves high accuracy throughout the genome, increasing linearly with coverage. The best algorithm’s sensitivity exceeded 80% for VAFs ≥4% and 15% for VAFs of 0.5-1% with 60x coverage.

The publicly available benchmarking data and comparative analysis of current methods provide a foundation for robust discovery of SV mosaicism in non-cancer tissues..

## Introduction

Somatic structural variants (SVs; ≥50bp genomic alterations) that emerge during human development^1^ give rise to genetic mosaicism, which is increasingly recognized as a major contributor to diseases not only in cancer but also in neurodevelopmental disorders^2,3^, neurological disorders^4^, and congenital anomalies^5^. Unlike somatic mutations in cancer, which are often identified by comparing tumor to matched normal tissue, somatic variants in non-cancer contexts may be present in multiple tissues, often without a clear “normal” comparator^1^. Detecting these events is challenging as their variant allele fractions (VAFs) are often below 5%, with signals that are easily overwhelmed by sequencing and mapping-induced artifacts, especially in repetitive or otherwise complex genomic regions^6,7^.

The Somatic Mosaicism across Human Tissues Network (SMaHT) is an NIH-funded program aimed at improving our understanding of the prevalence and impact of mosaicism across healthy human tissues^8^. Early results from this initiative have highlighted the complexity and challenges of identifying somatic mutations especially beyond single nucleotide substitutions^9^. Nevertheless, the undeniable importance of characterizing mosaicism necessitates improved methods to sequence, identify and compare variants accurately. This is especially true for SVs^10^, which are already challenging to detect in the germline and even more so in the somatic context due to their lower allele fractions and mosaic distribution^11^.

Recent studies have advanced variant benchmarking and accelerated genomics method development^12,13^. Multiple germline benchmarks exist (e.g., Genome in a Bottle (GIAB)^13,14^, Sequencing Quality Control 2 (SEQC2)^15,16^, Platinum Pedigree^17^), but few have addressed the complexity of somatic variants, with an exception of a GIAB study that reported 85 somatic single nucleotide variants (SNVs) for HG002^18^. Other benchmarking efforts have centered on tumor/normal somatic variant calling and provided a variant catalog for tumor-only SV^9,19^. However, these datasets do not capture the complexity of detecting mid- to low-VAF alleles and are often enriched for translocations, which are uncommon in non-tumour tissues^20,21^. Therefore, there is a need for mosaic benchmarks focused on SVs given the multiple reports highlighting the functional consequences of SV mosaicism^22,23,24^. Furthermore, as sequencing costs continue to drop, high coverage sequencing is becoming more affordable, making it easier to detect rare somatic mutations, especially if they are present across multiple tissues. To take full advantage of these improvements, we need well-characterized benchmark datasets that help develop and test methods for detecting variants present at low VAFs.

Here, we present a high-coverage, assembly-based benchmark that spans the full spectrum of somatic SV VAF (0.25-16.5%) with a sample derived by mixing six HapMap^25^ samples. From haplotype-resolved assemblies of each sample, we derived a benchmark for SV mosaicism detection containing 34,140 insertions and deletions ≥50 bp from across 90.2% of GRCh38 autosomes. The deep whole-genome sequencing data generated by the SMaHT Network for this sample include five Illumina replicates, four PacBio HiFi replicates, and two ONT replicates, with cumulative coverages of 1,745X, 380X, and 180X, respectively, which is sufficient to benchmark methods down to ultra-low VAFs. Using this resource, we systematically benchmarked 12 somatic SV calling strategies, providing insights into features that most impacted performance. To further aid interpretation of the results, we developed a novel approach for automatically assessing SV caller performance across SV stratifications. This benchmark, complemented by deep sequencing and orthogonal datasets (such as RNA sequencing and Element), provides a well-curated SV mosaicism benchmark with a complete germline and somatic SV catalog, coupled with high quality public data sets. Our analysis offers best-practice recommendations and the benchmark data provide a foundation for future bioinformatics efforts aimed at improving SV mosaicism detection.

## Results

### Design of the HapMap Mix sample

We generated the HapMap mix sample by blending genomic DNA from six well-characterised HapMap cell lines whose haplotype-resolved assemblies were produced by the Human Pangenome Reference Consortium (HPRC)^26^. This sample (**Figure 1a**) was built to create a broad spectrum of VAFs by including individuals at high abundance (HG005 at 83.5%), low abundance (HG02622 at 10%), and very low abundance (HG002, HG02257, HG02486 at 2% and HG00438 at 0.5%) thereby emulating VAFs of both germline and somatic variants. SV benchmark construction began with assembly-based SV calling on each of the six HPRC diploid assemblies, followed by cross-individual merging and SV harmonization to retain only SVs in high-confidence regions (**Figure 1b;** Methods). Each benchmark SV was annotated with its expected VAF, tandem-repeat context, and the number of neighboring SVs within ±500 bp, enabling fine-grained stratification of caller performance.

**Figure 1.**
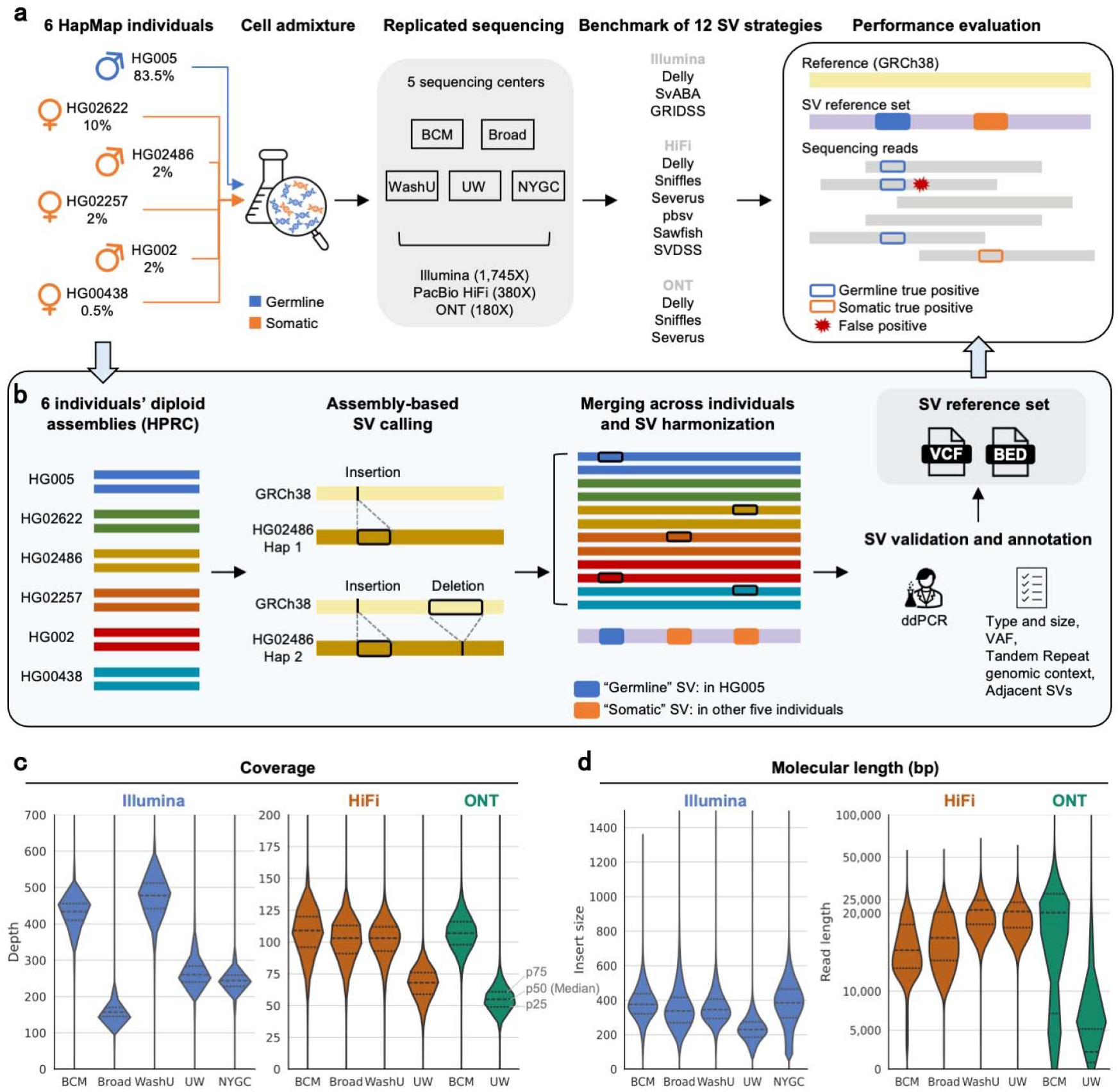
Experimental design of the SMaHT MIMS benchmark. **a)** Six HapMap samples with HPRC assemblies were in-vitro mixed at different ratios to represent germline and somatic variants. This sample was distributed across five Genome Characterization Centers (GCCs) and sequenced with short-read (Illumina) and long-read technologies (Oxford Nanopore, ONT and Pacific Biosciences, HiFi). Across sequencing runs, 12 somatic SV calling strategies were benchmarked against a high-confidence assembly-based SV set. **b)** The SV benchmark (Multiple Individuals in a Mixed Sample (MIMS)) was built using diploid assemblies from the HPRC. All SVs were merged and harmonized in a single VCF file and the variants were classified as germline if present in the high abundance sample (HG005, 83.5%) and somatic otherwise. Quality control of the mixes, validation and annotation was performed to select for high-confidence variants. **c)** Coverage statistics of all the WGS experiments by GCC and technology (based on coverage at one million randomly selected genomic bases). **d)** Read statistics of all the WGS experiments by GCC and technology.

To verify that the mixture ratios were maintained during library preparation, we quantified six representative SNV loci by droplet digital PCR (ddPCR), confirming design fractions within ±5% (**Supplementary Table 1**; Methods). The SMaHT HapMap Mix was then distributed to five SMaHT Genome Characterization Centers (GCCs) and sequenced with short-read sequencing (SRS) on Illumina and long-read sequencing (LRS) on Pacific Biosciences (HiFi) and Oxford Nanopore (ONT) platforms to obtain multiple replicates. All five GCCs generated a deep sequencing output, yielding 180-500X per replicate on Illumina, 70-110X on HiFi, and 65-115X on ONT (**Figure 1c**). Mean molecular length spanned 230-380 bp for Illumina, 16-22 kb for HiFi and 4-19 kb for ONT (**Figure 1d**). Additionally, four bulk RNA-seq replicates were produced by four GCCs, each with at least 150 million reads (**Supplementary Figure S1**), alongside sequencing experiments performed by individual GCCs (e.g., Element Biosciences sequencing at BCM). Tool and Technology Development (TTD) projects within the SMaHT Network produced additional datasets such as single-nucleus RNA sequencing (snRNA-seq) and ATAC-seq. All data sets are publicly available at https://data.smaht.org.

To validate the mixture ratios, we estimated VAFs at 51,428 tandem repeat (TR) loci^14^ in each LRS replicate (Methods). For each TR locus, the length of each aligned read spanning the region was assigned as support for the closest allele length among the HPRC assembly haplotypes aligned over the same region (Methods). This process allowed calculation of the observed VAF. The standard deviation of the observed minus expected VAF difference was 0.07 for HiFi and 0.11 for ONT (**Figure 2a; Supplementary Figures S2a**,**b**). Regression of observed vs. expected VAF gave slopes of 0.92 with an intercept of 0.02 and Pearson correlation of 0.94 for HiFi, and a slope of 0.85 with an intercept of 0.05 and correlation of 0.90 for ONT (**Supplementary Figure S2c**). These results were in line with previously reported patterns of VAF variance^27^ and confirmed that each sequencing experiment was consistent to the expected proportions of HapMap alleles.

**Figure 2.**
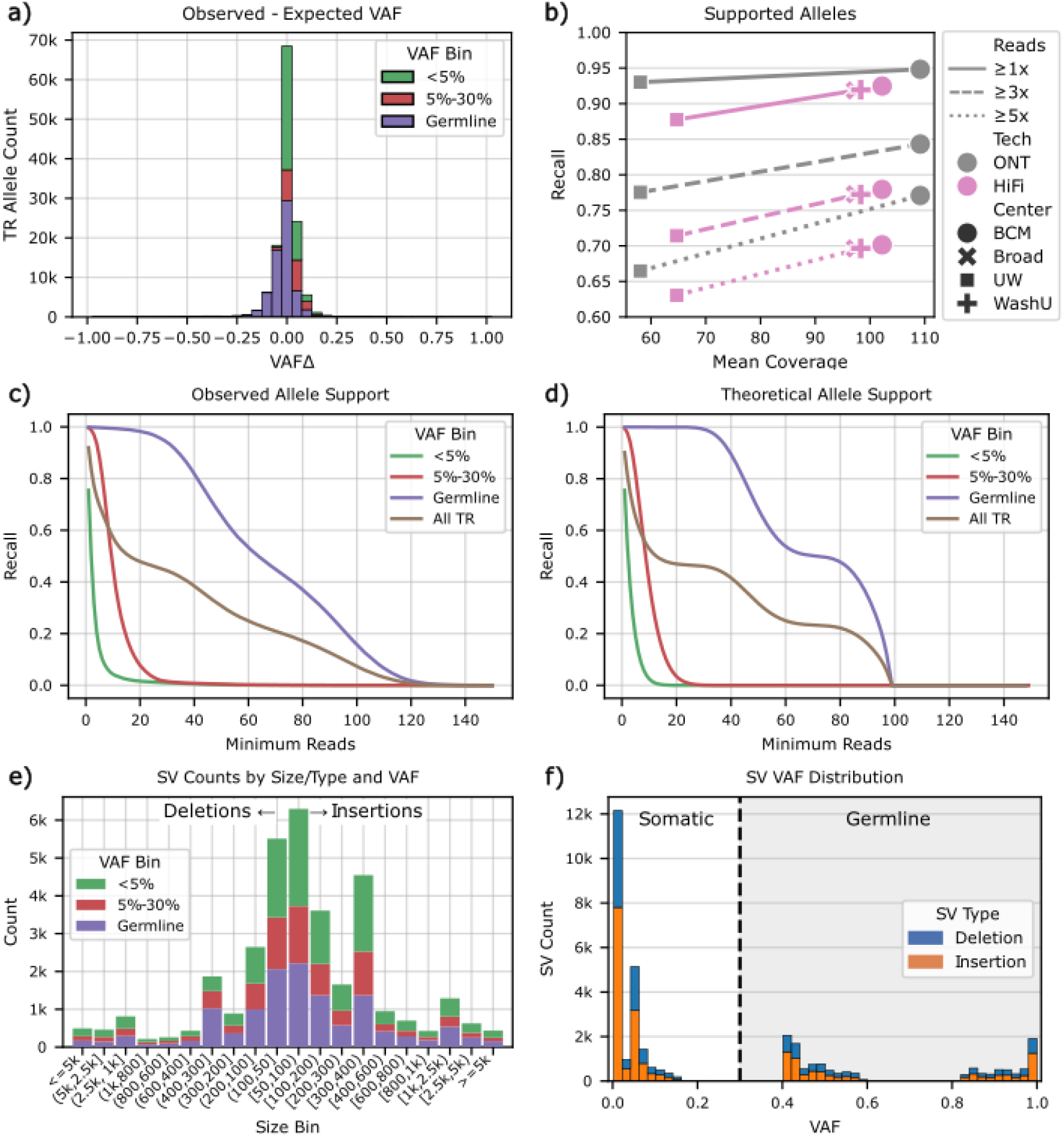
Sequencing Validation and Benchmark SV Composition. **a)** Observed minus expected VAF over TR alleles for a 100x HiFi sequencing experiment. Histogram binwidth 0.05. Hue is expected VAF Bin. All sequencing experiments available in Supplementary Figure 2b. **b)** Recall for TRs with minimum read support of 1x, 3x, and 5x for six sequencing experiments using two long-read technologies from four sequencing centers. Note that Broad and WashU HiFi experiments overlap. **c)** Observed Cumulative Distribution Function (CDF) of recall on 100x HiFi sequencing. All sequencing experiments plotted in Supplementary Figure 2c. **d)** Theoretical CDF of the beta-binomial process modeled on 100x coverage. **e)** SV Count by Size, Type, and Variant Allele Fraction (VAF). **f)** SV VAF Distribution for deletions and insertions; Histogram binwidth 0.02.

In the TR allele length analysis, alleles having read support (i.e., recall) decreased with increasing minimum support thresholds, from a mean of 0.91 at ≥1x to 0.77 at ≥3x and 0.69 at ≥5x (**Figure 2b**). More broadly, the cumulative distribution of TR allele recall under varying support thresholds depended on the expected VAF and sequencing depth. **Figure 2c** shows results from a representative 100x HiFi sequencing experiment; all experiments are shown in **Supplementary Figure S2d**. These distributions illustrate that low-fraction alleles (e.g., 1% VAF) are less likely to meet moderate support thresholds (e.g., ≥2 reads) at moderately high sequencing depths (e.g., 100x), leading to reduced recall. To generalize this behavior and enable predictive estimation of recall, we modeled the probability of observing sufficient read coverage as a beta-binomial process (Methods; **Figure 2d; Supplementary Figure S2e**). For the representative 100x HiFi sequencing experiment, our beta-binomial model achieved a log-likelihood of −422,379 compared to −618,820 under the standard binomial model, indicating a substantially better fit to the observed data. This improvement reflects the beta-binomial’s ability to account for overdispersion in read support counts, and was found to be consistent to all long-read sequencing experiments (**Supplementary Table 2**). This modeling analytically relates sequencing depth, VAF, and read support thresholds to the probability of observing sufficient evidence for detection, enabling estimation of the maximum achievable recall of somatic alleles under different experimental designs.

### Mosaic Benchmark SV Creation

The haplotype-resolved assemblies from HPRC^26^ were used to identify high-confidence SVs (50 bp to 50 kbp) and define reliable genomic regions that comprise the SMaHT Multiple-Individuals in a Mixed Sample (MIMS) SV benchmark. Each haplotype was aligned independently with minimap2 and variants called with paftools^28^, resulting in a consolidated set of 54,291 SVs (37,056 insertions; 17,235 deletions) within the high confidence regions (described below). Independent alignment of haplotypes gives rise to redundant SVs where identical alleles appear different due to e.g., alignment ambiguities^29,30^. Typically, SV merging approaches are used to consolidate putatively redundant SVs^31,32^. However, these processes rely on heuristic similarity thresholds to identify redundancies. This is a lossy operation that is subject to overmerging, a phenomenon where highly similar yet distinct alleles are misidentified as being the same^31^. To create the full set of distinct, non-redundant alleles, SV harmonization was performed with truvari phab^14^. This tool reconstructs original haplotypes over SV regions of the genome before performing a multiple-sequence alignment that produces a more parsimonious set of SVs. This process reduced the SV count by 10.4% to 47,638 (33,390 insertions; 14,248 deletions). Details on SV counts from the original alignments, harmonized alignments, and a comparison to classic SV merging can be found in **Supplementary Section 2**.

With all haplotypes’ SVs harmonized, all remaining SVs are distinct. Importantly, distinct SVs can differ by as little as 1 bp. For example, two insertions at the same position and of the same length may differ internally by a single SNP, resulting in two distinct SVs. In total, 8,813 SVs (18.5%) contain only substitution mutation differences to other SVs with equal type, position, and length. Moreover, 18,119 SVs (38%) have equal type, position, and are within 5bp of length to another SV. These minor differences between SVs present significant challenges to SV benchmarking. First, SV callers are designed to detect larger structural differences between alleles and often collapse alleles with minor discrepancies to compensate for sequencing errors or alignment artifacts^10^. Second, small errors in reported SVs, such as a slight misestimate in the length of a germline SV, can mistakenly cause it to be best matched with a distinct somatic SV, leading to unreliable measurement of a tool’s performance.

To address the challenges presented by minor differences between SVs, we construct a primary benchmark VCF by collapsing the harmonized SVs with high similarity using dynamic thresholding. This process restricts merging to SVs of the same type within 100 bp of each other and uses a capped size similarity threshold to collapse only SVs with between 5 bp and 30 bp length differences (Methods). This produced a final set of 34,140 SVs (20,545 insertions; 13,595 deletions; **Figure 2e**). SVs were annotated with expected VAFs derived from the expected proportions of samples in the mix and calculated as the weighted sum of haplotypes in which the variant occurred (**Figure 2f**). VAFs in the range of germline heterozygous or homozygous variants (41.75% - 100%) were achieved on SVs present in at least HG005 and comprised 12,756 (37.3%) SVs with the remaining 62.7% having low VAF (0.25% - 16.5%) deemed somatic. SVs were annotated by their proximity to tandem repeats (TRs; 57.7% of SVs) and the number of neighboring SVs within 1 kbp (36.3% ≥1; **Supplementary Figure S4**). These annotations (SV type, SV size, expected VAF, tandem repeat context, and neighboring SVs) provided stratifications for more granular benchmarking.

To create high confidence regions for the benchmark, all assembly haplotypes’ alignments were required to produce 1x coverage for a region. The intersection of all haplotypes’ confidently covered regions spanned 91.1% (2,617,765,246 bp) of GRCh38 autosomes. Long tandem repeats (≥10 kbp) or segmental duplications not fully spanned by the confident regions, and regions within 5 kbp of a reference gap were also excluded. Finally, benchmark SVs were clustered to all neighboring SVs within 1 kbp to identify if any one haplotype produced two or more SVs. If so, the entire region was deemed as having high density/complexity and excluded from the benchmark regions. These exclusion criteria produced 9,774 benchmark regions spanning 90.2% (2,594,105,129 bp) of GRCh38 autosomes.

A subset of SV calls were orthogonally validated using droplet digital PCR (ddPCR) to precisely quantify SV breakpoint sequences within the sample mixture. A random selection of somatic SVs (i.e., not present in HG005) were chosen and ddPCR assays were designed on four SVs, with all validating and having an observed VAF near the expected with an average absolute delta of 1.09 (**Supplementary Figure S5, Supplementary Table 3**).

### Caller evaluation across stratified challenge categories

We benchmarked eight SV callers capable of detecting mosaicism (Delly^33^, GRIDSS^34^, pbsv^35^, Sawfish^36^, Severus^37^, Sniffles^11^, SvABA^38^, SVDSS^39^). SV callers were chosen based on their operation without a matched control sample and ability to report SVs at VAFs <10%. Each caller was run on up to three sequencing technologies (Illumina, HiFi, and ONT), producing 12 SV discovery pipeline results (**Figure 1a**, right; Methods). With the availability of sequencing replicates across GCCs, a total of 45 SV discovery results were obtained. Among the evaluated tools, only Sniffles separately reports germline and somatic SVs whereas all others return an undifferentiated single set of SVs. To score callers fairly, we intersected each call set to the SMaHT MIMS benchmark using Truvari^31^ (Methods).

To assess the performance of somatic SVs, precision and recall was calculated by excluding germline benchmark SVs and callers’ false positives with a reported VAF above 25%, thus isolating analysis to each algorithm’s ability to detect low-VAF events. Across 45 call sets, recall ranged from 4-50%, and precision from 36-93% (**Supplementary Figure S6**), underscoring the difficulty posed by the MIMS benchmark. After binning the benchmark set by expected VAF (<0.5%, 0.5-1%, 1-4%, 4-5%, 5-10%, 10-20%; **Supplementary Figure S7**), VAF-bin-specific recall revealed persistent gaps: even in the deepest replicate within 10-20% bin, the best caller per platform reached only 27% (Illumina), 88% (HiFi), and 82% (ONT) (**Figure 3a**). These shortfalls implicate challenges beyond coverage and VAF.

**Figure 3.**
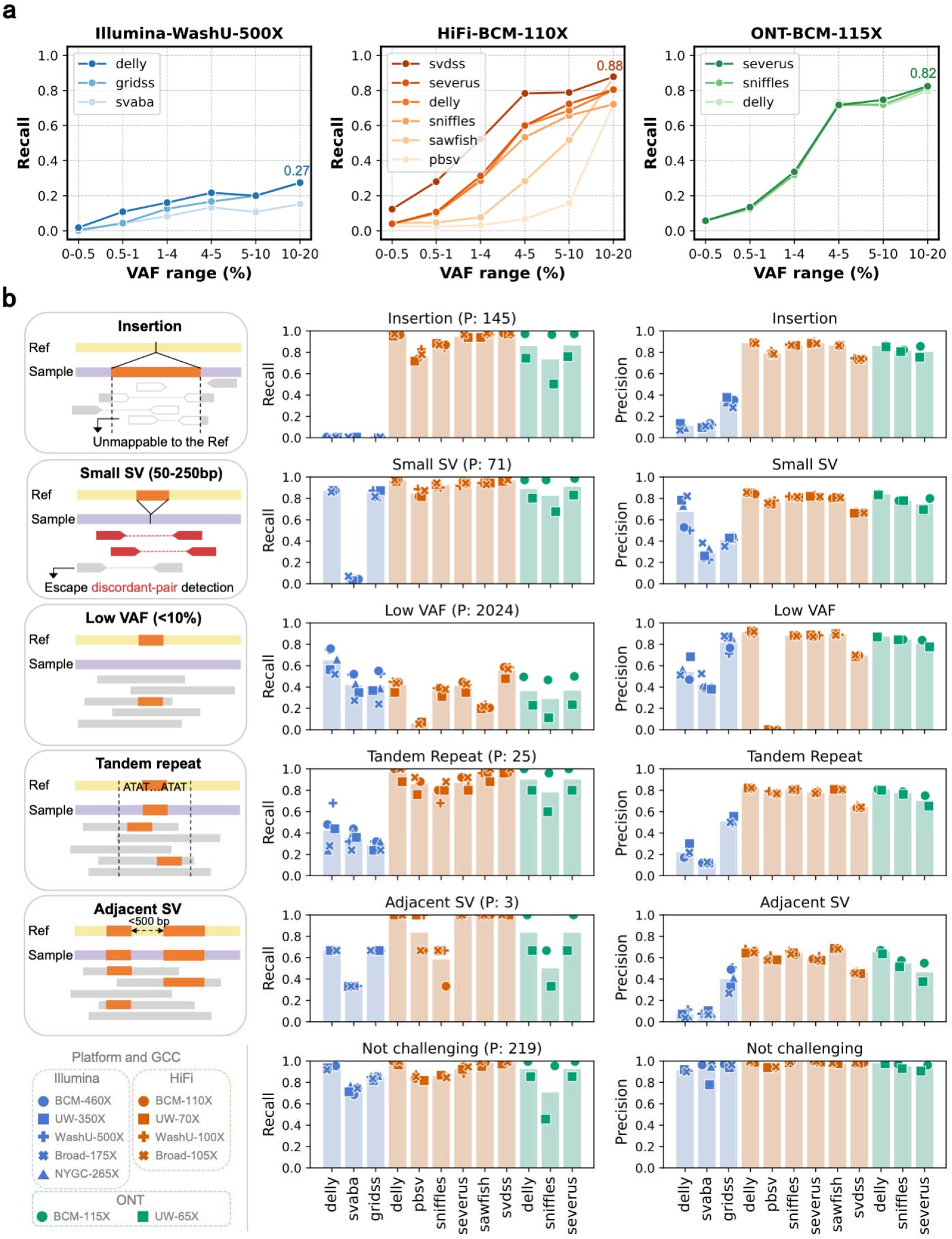
Benchmark 12 somatic SV calling strategies. **a)** Recall by VAF bin for each platform (Illumina, HiFi, and ONT), shown for the replicate with the highest sequencing depth. Callers are ordered by overall recall; darker color indicates a higher recall. VAF bins by expected VAF are <0.5%, 0.5-1%, 1-4%, 4-5%, 5-10%, 10-20%. **b)** Difficulty-stratified performance. Left: each SV is assigned to one difficulty category (insertions, small SVs (≤250 bp), low VAF, tandem-repeat overlap, or clustered/adjacent SV) or to a residual “not- challenging” class. Middle & right: recall (middle) and precision (right) are reported for 12 workflows. Recall is calculated on difficulty-exclusive subsets, with sample size P (the number of calls in the subset) shown in headers. The bottom row (“Not challenging”) contains variants lacking any annotated difficulty. Dots show results from individual sequencing replicates (marker shape and color represent replicate and platform, separately, with depth highlighted in the legend), and vertical bars denote the replicate mean for each workflow.

To dissect the underperformance, we annotated every SV with five features found to obscure detections: (1) SV type (insertion versus deletion), (2) small size (50-250 bp), (3) low VAF (<10%), (4) overlap with tandem repeats, and (5) proximity to another SV’s breakpoint within ±500 bp, capturing multi-allelic or adjacent events. Because these features often co-occur and low-VAF <10% events constitute 94% of the benchmark’s somatic SVs, we calculated recall on mutually exclusive difficulty subsets (**Supplementary Figure S8**). The resulting subsets contained 145 insertions, 71 small SVs, 2024 low-VAF SVs, 25 tandem-repeat SVs, 3 adjacent SVs, and 219 SVs that are outside of any difficulty above (“not-challenging”).

Caller and platform performance diverged largely across these difficulty strata (**Figure 3b**). In the “not-challenging” set, comprising 1% of the somatic baseline SVs, most workflows exceeded 80% recall and precision, confirming that our stratification represents the key obstacles faced by every algorithm. Insertions were the clearest fault line between LRS and SRS. LRS approaches maintained comparable recall to the not-challenging set, whereas all SRS callers missed most insertions: Illumina-based mapping handles inserted sequences poorly; on average, SRS callers reported 1.4k insertions versus 11k for LRS callers (**Supplementary Figure S9a**). SRS approaches captured insertions only if they are shorter than the read length (often <100 bp), exhibiting high precision but low sensitivity; insertions exceeding the read length were usually tandem duplications inferred from clipped reads, containing many false positives and driving down precision for the insertion category (**Supplementary Figure S9b**).

Small SVs posed a second major challenge to short reads. They often cause only subtle insert-size shifts, leading supporting reads to be misclassified as concordant pairs and making it difficult to distinguish real SVs from mapping artifacts. Among the three SRS callers, SvABA detected the fewest small events, missing most deletions within the alignment of a single read (**Supplementary Figure S9c**). Delly and GRIDSS captured small SVs at rates comparable to LRS callers but at the cost of reduced precision, reflecting an enrichment of false positives across the small size range (**Supplementary Figure S9b**,**d**).

Detection of low-VAF events requires both deep sequencing and effective filtering of random artifacts. Delly on SRS achieved the highest recall because the Illumina data were sequenced at greater depth, and per-replicate recall rose monotonically with coverages. In contrast, two HiFi-specific callers, pbsv and Sawfish, struggled as pbsv recovered a few calls below 10% and for Sawfish below 6% VAF (**Supplementary Figure S10a**). This might be because of their consensus approaches. Among SRS callers, only GRIDSS effectively controlled low-VAF false positives; precision for Delly and SvABA showed a surge of false positives for VAF<4% (**Supplementary Figure S10b**). Most LRS callers retained high precision across the depths; SVDSS was the exception, which exhibited a similar spike in false positives below 3% VAF (**Supplementary Figure S10c**). Notably, precision for LRS callers was largely depth-independent, indicating robust filtering of stochastic noise, whereas SRS precision sometimes declined at the highest coverages as additional artifacts were admitted.

Tandem repeat loci and adjacent SV breakpoints further complicated SV detection and representation. SVs within tandem repeats suffered substantial recall and precision losses for all SRS callers due to mapping ambiguity. Adjacent SVs reduced precision for every workflow, often because neighbouring signals were mis-merged or mis-placed (**Supplementary Figure S11**).

Overall, the stratification confirms that insertions and tandem-repeat regions remain blind spots for SRS, while highlighting caller-specific strengths and weaknesses that will guide future algorithmic improvements.

### Evaluation of platform-specific detection limits

The previous result section highlighted that a driving factor of recall in this MIMS study is the overall coverage. Higher coverage should enable higher read support and thus also the identification of ultra-low VAF SV. To assess this, we merged the different replicates across the GCCs to evaluate the performance constringent on the coverage levels. Adjacent benchmark SVs were excluded to avoid distorting allele fraction. For each platform we calculated VAF-bin-specific recall for all 12 calling strategies and visualized the two best performers per platform as representative curves (**Figure 4**). We estimated an “upper-bound” recall obtained by running a platform-specific genotyper and counted the true calls that are supported by 1, 2, or 3 supporting reads (Methods).

**Figure 4.**
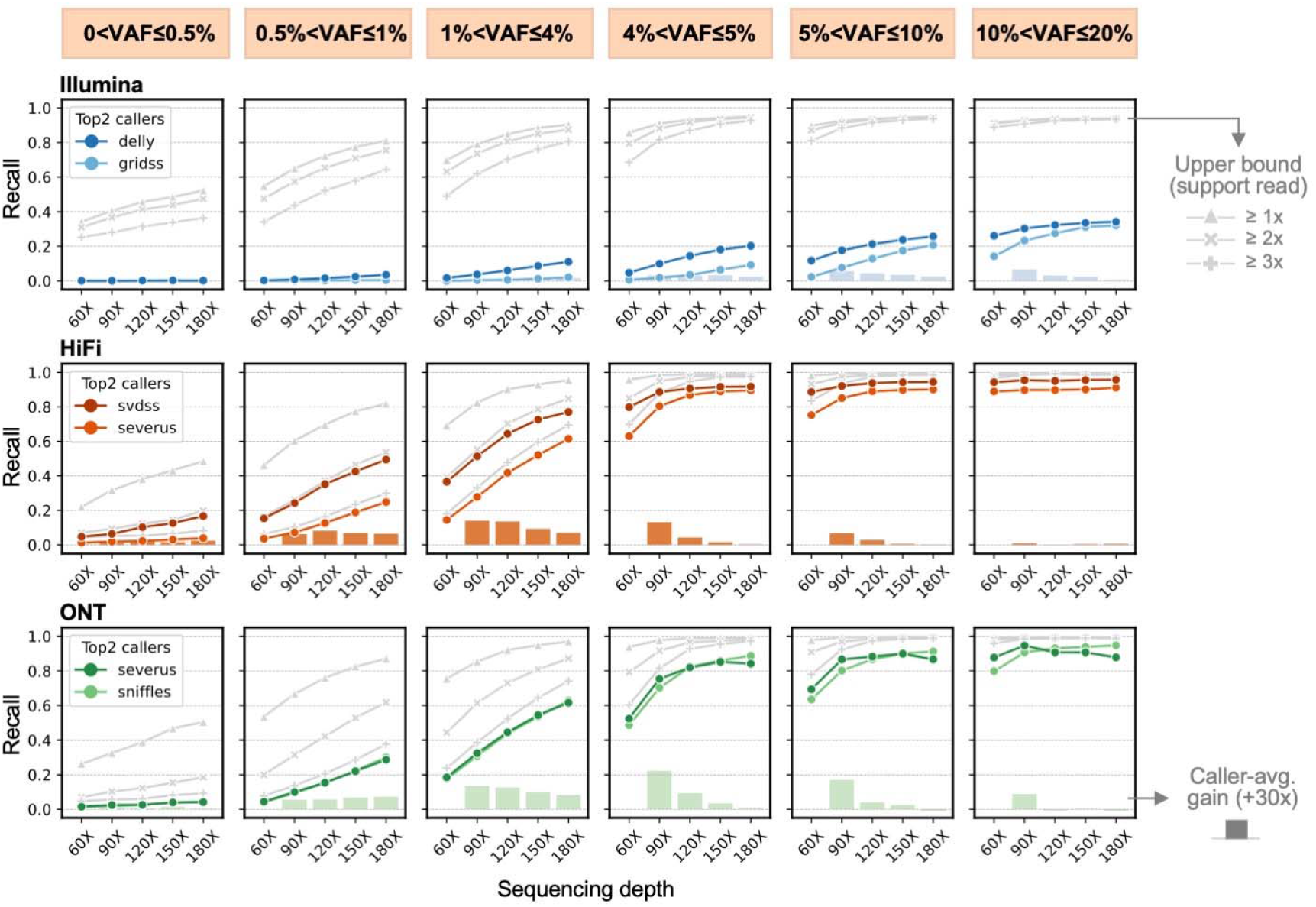
Platform-specific detection limits for somatic SVs. Recall across VAF bins at fixed sequencing depths. For each platform (Illumina, HiFi, ONT), the two best-performing callers are plotted; each dot marks the recall achieved in a given VAF bin. Grey lines show the genotyper-estimated recall if detection requires 1, 2, or 3 supporting reads for each sample. Grey vertical bars indicate the average recall gain (averaged across two callers) for a 30x increase in coverage relative to the depth immediately to the left, illustrating the marginal benefit of additional sequencing.

Illumina recall rises with depth but interestingly plateaus at 35% even at 180x coverage, which is well below the estimated recall. This indicates that limited read length rather than coverage is the main constraint. Stratified analysis reveals that the plateau is driven by three sources of reduced sensitivity: insertions, small SVs, and variants overlapping tandem repeats. When the analysis is restricted to large deletions in non-repetitive (“easy-to-map”) regions, Illumina recall increases to 90% at 120x coverage (**Supplementary Figure S12**). Despite this observation SRS remains more practical to achieve this coverage level given the associated costs.

LRS delivers far greater sensitivity than SRS: at 60x HiFi, SVDSS exceeds 80% recall for VAFs ≥4%, approaches 40% at VAFs 1-4%, and captures ∼10% even for VAFs down to 0.5%. While SVDSS is tailored to accurate HiFi reads, results from Severus,which runs on both HiFi and ONT, show that the two LRS platforms achieve similar recall at the same depth. The detection limit of LRS exhibits an almost linear rise with coverage, although the marginal gain depended on VAF. Bars in **Figure 4** quantify the mean recall gain per 30x increment (average across the two representative callers on each platform). In the 0.5-1% VAF bin, successive 30x increments contributed nearly identical gains, whereas above 4% the gain decreased and ultimately disappeared, signalling saturation. Observed performance aligned closely with the estimated upper bounds: SVDSS reports an SV if two supporting reads are present, while Severus and Sniffles used three reads as default threshold.

VAF-bin-specific precisions stabilized at ∼80% for VAF>1% events once depth reached 60X for LRS and 120x for SRS. LRS approaches sustained this level across all VAF bins, except SVDSS, which drops by 10% in the 1-4% bin owing to an influx of low-VAF false positives. With SRS, GRIDSS approached 80% precision at VAF 1-4% when depth reached 120x, whereas for VAF>4% the same precision was achieved at 90x (**Supplementary Figure S13**).

### Toward Standardized Variant Caller Assessment

The identified patterns in pipeline performance required manual curation, a separation of germline and somatic alleles, and the use of subsets of the benchmark. These bespoke processes, while insightful to understanding detection of SV mosaicism, ultimately under-sample the number of loci available in the benchmark and are not easily scalable for broader adoption. To remedy this, we developed a statistical test to automate the identification of stratifications that impact a result’s accuracy. This “StratP” test enables statistically rigorous prioritization of stratifications by testing the null hypothesis that a result’s accuracy is independent of a given stratification (Methods). We provide the StratP test as a new sub-command of Truvari. StratP operates on a single Truvari benchmark result and produces a table of a caller’s accuracy (recall or precision) on each stratification along with a p-value which, if below the significance threshold, indicates a dependency between a caller’s performance and the stratification.

Left-tailed StratP tests with significance threshold set to 0.01 were performed on the 45 SV benchmarking results (i.e., combinations of SV callers, technologies, and GCCs) and replicated the findings of manual curation (**Supplementary Table 4**; **Supplementary Section 5**). To summarize, across all results and stratifications, the average recall was 0.412 with germline stratifications averaging 0.667 and somatic 0.267. The StratP test identified seven stratifications that significantly reduced recall of at least one result and were therefore considered difficult (**Table 1**). Difficult stratifications’ average recall was 0.319 compared to 0.462 for the rest (**Supplementary Figure S14**). Low VAF (<5%) baseline SVs significantly reduced recall for 39 of the 45 results and averaged 0.101 recall. The 6 results not impacted by low VAF (<5%) were all high-coverage (≥246x) Illumina based results from Delly and SvABA. Five of the difficult stratifications were specific to short or long reads: four were associated with Illumina-based callers (Insertions, TR regions, <200bp length), while one (somatic VAF 5%-30%) was exclusive to the germline-focused long-read callers pbsv and Sawfish.

**Table 1.** **StratP Test Results** of stratifications identified as significantly (alpha=0.01) impacting a result’s recall (Reject null-hypothesis), compared to the number of results where the stratification did not significantly impact recall (Accept null-hypothesis). Tallies are of 45 SV benchmarking results from 8 SV callers applied to 3 sequencing technologies from 5 GCCs. Total results per-technology were Illumina 15, HiFi 24, ONT 6. Bold/highlighted cells are stratifications with Reject/Accept results unique to short or long-read (HiFi & ONT) sequencing.

| Stratification | SV Count | Reject $H_0$ Results | | | | Accept $H_0$ Results | | | |
| --- | --- | --- | --- | --- | --- | --- | --- | --- | --- |
|  |  | Total | Illumina | HiFi | ONT | Total | Illumina | HiFi | ONT |
| Somatic (VAF $< 5\%$) | 13,184 | 39 | 9 | 24 | 6 | 6 | 6 | 0 | 0 |
| Neighboring | 12,401 | 35 | 13 | 17 | 5 | 10 | 2 | 7 | 1 |
| Insertions | 20,545 | 15 | 15 | 0 | 0 | 30 | 0 | 24 | 6 |
| TR Regions | 19,712 | 13 | 13 | 0 | 0 | 32 | 2 | 24 | 6 |
| Small [100,200) bp | 6,256 | 9 | 9 | 0 | 0 | 36 | 6 | 24 | 6 |
| Somatic (VAF 5%-30%) | 8,200 | 7 | 0 | 4 | 3 | 38 | 15 | 20 | 3 |
| Small<br>[50,100) bp | 11,810 | 5 | **5** | 0 | 0 | 40 | 10 | 24 | 6 |

These results confirm that short-read callers perform well on somatic SVs, but with the caveat that the SVs are isolated deletions outside of tandem repeats, which account for 30.8% of the benchmark. Furthermore, long-read callers can perform well on somatic 5%-30% VAF SVs, but struggle to recover SVs below 5% VAF relative to their performance on other stratifications. Regardless, the ability of long-read callers to discover insertions and SVs within tandem repeats produced a higher overall recall of somatic SVs than short-reads. For somatic SVs, the top mosaicism detecting long-read callers (SVDSS, Severus, Delly, Sniffles) had average recall of 0.35 compared to the top short-read callers (Delly, GRIDSS) averaging 0.10. More generally, these analyses demonstrate the StratP test as a powerful tool for SV benchmarking and exploratory data analysis, enabling easy identification of variant features that most affect caller performance.

### Validation with Orthogonal Technologies

We used additional sequencing experiments utilizing different platforms or molecules. The BCM GCC performed a WGS experiment using the Element Bioscience short-read Aviti platform, producing 232x genome coverage. We replicated the Illumina analysis pipelines of Delly, SvABA and GRIDSS, as well as directly genotyped deletions with SVTyper^40^ (**Figure 5a**). The Aviti platform achieved similar performance to the Illumina sequencing, with Delly showing the highest recall at 37.65% and SvABA the highest precision at 76.04% (**Supplementary Figure S15**). SVTyper’s genotyping recall of all deletions was 51.72% and 70.37% outside of TR regions. Further stratification of the genotyping results by VAF showed a 14 percentage point higher germline recall than somatic. Notably, the VAF reported by SVTyper was largely lower than the benchmark’s expected VAF (**Supplementary Figure S16**).

**Figure 5.**
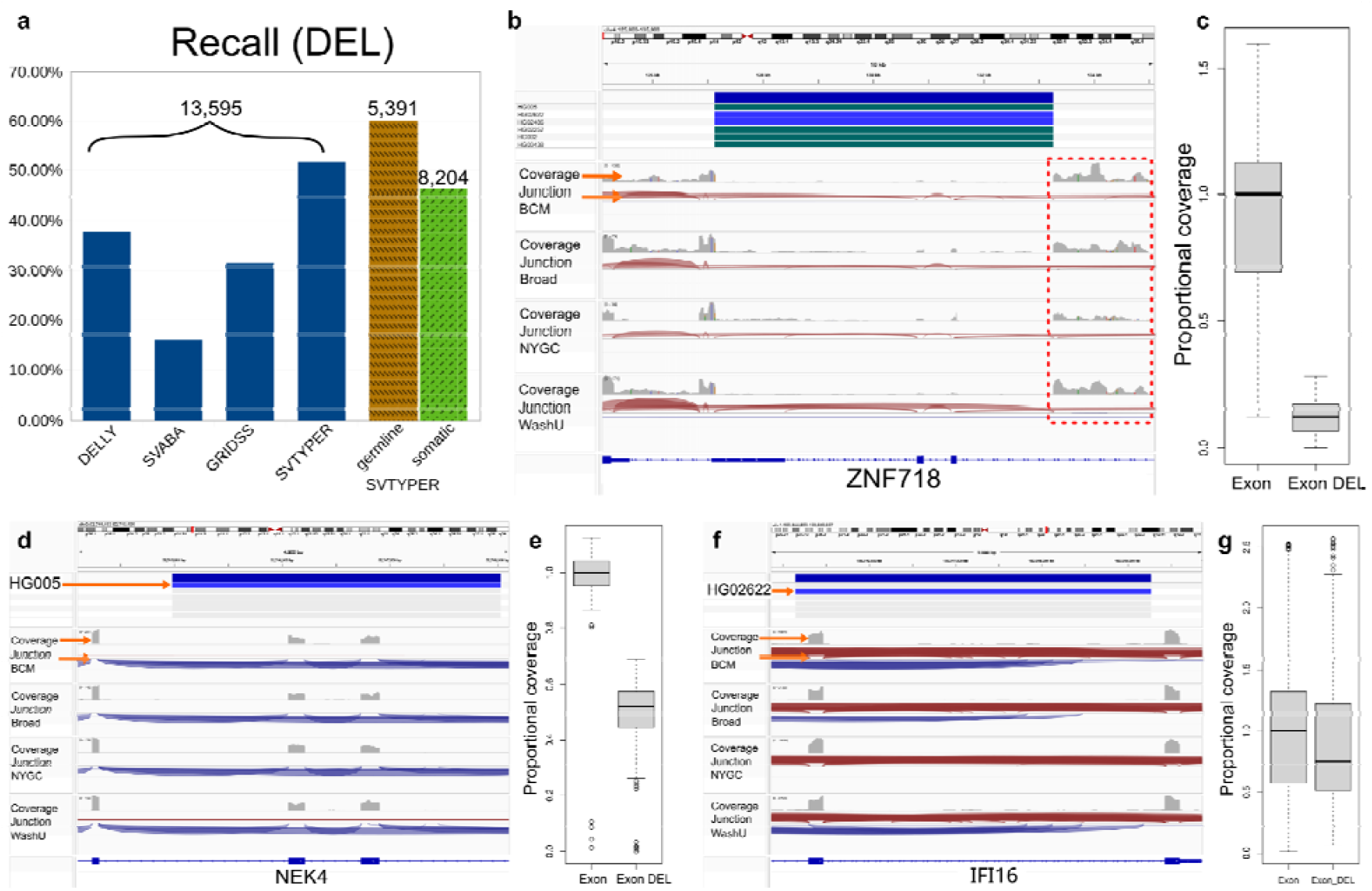
Validation of orthogonal methods with the SMaHT MIMS benchmark. **a)** SV recall from a WGS experiment using the Element Bioscience short-read Aviti platform (232x coverage) used as orthogonal validation. Recall of DEL from three short-read SV callers and genotyper are shown. For the genotyper we stratified the SV based on VAF. The numbers shown are the total number DEL to be detected and proportion of detection is shown in the y-axis. **b)** IGV screenshot of a germline DEL (expected VAF ∼94%) that overlaps with three exons of the ZNF718 gene. Shown (from top to bottom) are the benchmark SV followed by coverage, junction information (marked with orange arrows) and reads tracks for the four replicate samples: Baylor College of Medicine (BCM), Broad Institute Inc (Broad), New York Genome Center (NYGC) and Washington University (WashU). **c)** Boxplot of the coverage distribution of the three exons overlapping and not overlapping the benchmark SV shown in panel b. **d)** IGV screenshot of a germline DEL (expected VAF ∼42%) that overlaps with two exons of the NEK4 gene. Both affected exons show lowered coverage similar to the expectation based on the sample mix and genotype (observed 55.4% and 58.1% coverage, expected 58%). **e)** Boxplot of the coverage distribution of the three exons overlapping and not overlapping the benchmark SV shown in panel d. **f)** IGV screenshot of a somatic DEL (expected VAF ∼5%, highlighted with an arrow) that overlaps with an exon of the IFI16 gene. The affected exon shows lowered coverage slightly higher to the expectation based on the sample mix and genotype (observed 25% and expected 5%). **g)** Boxplot of the coverage distribution of the two exons overlapping and not overlapping the benchmark SV shown in panel f.

We further analyzed four replicates of bulk RNA-seq, assessing the potential of the functional and variant calling within one assay. The BCM, Broad, NYGC and WashU GCCs reported 201.4, 175, 377, and 288.1 million reads, respectively (**Supplementary Figure S1**). Since no RNA-seq SV callers currently exist, gene annotation was used to identify benchmark SVs with RNA-seq support that overlapped expressed, protein coding genes. Expressed genes were found using two normalization methods widely used in RNA-seq that account for both sequencing depth and gene length. Transcripts Per Kilobase Million (TPM) normalizes reads counts by gene length in kilobase to then scale it with a million factor, while Fragments Per Kilobase Million (FPKM) normalizes read count by gene length and the total number of mapped reads in a sample. For deletions, we selected SVs that overlapped with at least 50% of any exon of an expressed gene (Methods). Over all four replicates we found 539 germline SV-gene overlap candidates and 213 somatic SV-gene overlap candidates in the benchmark. **Figure 5b** shows a deletion with an expected VAF of 94%, that affects three exons of the *ZNF718* gene (one partial, two full). Coverage drops can be observed at a slightly lower than expected proportion based on the mix (observed 88.71% and 84.58%, expected 94%, **Figure 5c**). Furthermore, we observe coverage over the intron after the deletion, a sign of intron retention. Another example worth highlighting is a germline deletion with an expected VAF of 41.75% that overlaps with two exons of the *NEK4* gene (**Figure 5c**). Both exons overlapping with the deletion show a drop in coverage of 44.6% and 41.9% respectively (**Figure 5d**). For somatic variants, we detected a deletion with an expected VAF of 5%, overlapping one exon of the *IFI16* gene (**Figure 5e**). Coverage drop can be observed at a higher than expected proportion based on the mix (25% and 5% respectively, **Figure 5f**). Additional examples of deletions overlapping exons at germline and somatic levels were found that were supported by lowered coverage and skipped exons based on junction information (**Supplementary Figures S17, S18**). Further, we detected a split read result of a deletion in an intron of the *GDAP1* gene (**Supplementary Figure S19**).

For insertions, we also performed expressed gene and SV overlap. **Supplementary Figure S20** shows an insertion with an expected VAF of 100% that overlaps an intron of *IL19*. We observed clip reads in all the read overlaps with the SV. Additional examples are shown in **Supplementary Figure S21**, which depicts an insertion with an expected VAF of 44%, that disrupts transcription of the *MAN1B1* gene shown as a drop in coverage in the exon; **Supplementary Figures S22, S23** show and insertion overlapping with exons of the *CCDC92* and *PLA2G4C* genes. In both cases, we observe soft-clip reads and increase in coverage that resemble a duplication. Thus overall multiple SVs in the benchmark had support within RNA-seq. We hope that this resource will also spark interest in the community to develop RNA-seq based SV callers capable of detecting germline or somatic events.

## Discussion

This study introduces the first benchmark specifically designed to evaluate the detection of structural variant (SV) mosaicism, a long-standing gap in the somatic variant analysis space. By creating a six-sample mixture with controlled proportions, we captured SVs across defined variant allele fraction (VAF) bins, enabling rigorous performance stratification across the somatic detection spectrum. While this experimental design introduces an inflated number of apparent somatic SVs due to admixture artifacts, we addressed this limitation through careful curation, merging of near-identical calls, and stratification of SVs across multiple variant properties to produce a high-confidence, VAF-aware benchmark. Despite its synthetic origin, the benchmark offers a reproducible and quantitative framework for assessing SV mosaicism detection methods under realistic and tunable conditions. This is especially timely, as we are at a pivotal moment in mosaicism research. Technical advances, falling sequencing costs, and deeper biological insights have brought the field to the cusp of routine somatic SV analysis. Yet, major challenges remain, particularly in achieving reliable detection of low VAF variants, enabling interpretation across variant types, and incorporating mosaicism into clinical genomics pipelines. This benchmark lays a foundation for the field to tackle those challenges. By providing an open, VAF-aware resource with both genomic and transcriptomic resources, we aim to catalyze a new generation of tools and studies that will move SV mosaicism from a niche topic to a core component of precision medicine and human disease research.

Our results reinforce that sequencing coverage remains a key driver of somatic SV detection, particularly for low-VAF variants. In bulk sequencing, increasing depth improves sensitivity, but only to a point. Beyond a certain coverage threshold, diminishing returns and practical limitations (cost, DNA input, tissue availability) constrain the utility of ultra-deep sequencing. Interestingly, we found that long-read sequencing platforms exhibit a more consistent and predictable increase in recall across VAF bins than short-read sequencing despite the lower coverage available. Through extensive manual curation and the design of a new statistical test of variant stratification performance, we found this was explained by long-reads enabling better recovery of insertions and repeat-associated SVs. In contrast, Illumina-based methods showed a coverage plateau with limited gains, suggesting that technical blind spots, not coverage, may cap detection performance. These results emphasize that while long-read technologies are currently the most promising path forward for sensitive and unbiased somatic SV detection, cost and DNA requirements still pose major barriers to broad adoption, particularly for studies involving rare tissues or large cohorts.

The goal of this SMaHT MIMS SV benchmark is to support the development of somatic SV detection tools by providing both a practical and aspirational target for performance evaluation. The primary benchmark collapsed similar SV alleles at a locus when they differed by less than 5-30 bp to ensure remaining alleles are separated by differences both detectable by current pipelines and likely to be biologically meaningful. This allows tools to report ‘overmerged’ SVs without being penalized for missing near-duplicate alleles, while still capturing informative distinctions between somatic SVs. Although this lower-resolution benchmark reflects the current limitations of SV detection, it provides a realistic and useful standard for near-term tool development. We also make available the harmonized VCF, which preserves the full complexity of allele diversity observed across the 12 HapMap-derived haplotypes, including SVs that differ by as little as a single base pair. This high-resolution benchmark is designed to drive tool development toward resolving fine-scale somatic variation, such as low-VAF SNPs internal to germline SVs. Tools that succeed against this more stringent benchmark will help pioneer a new era of somatic and mosaic variant discovery, deepening our understanding of human health and disease.

The SMaHT Network is uniquely positioned to lead this transformation. Through its coordinated efforts across technologies, tissues, and analytical frameworks. SMaHT is generating the next generation of reference materials for somatic and mosaic mutation detection. Resources such as this benchmark will be critical for driving innovation and establishing standards needed for translational and clinical integration. To facilitate progress across the field, we have made this benchmark publicly available and designed it to be extensible, with complementary layers of RNA sequencing data. This enables benchmarking of variant detection as well as exploration of the downstream impact of somatic SVs on gene expression and splicing. However, no current tools exist to detect somatic SVs from RNA-seq data, representing a key unmet need. We hope that this resource inspires the development of novel methods, both computational and experimental, that bridge this gap and enable functional insights into somatic variation within the same assay.

## Methods

### Sequencing

The Somatic Mosaicism across Human Tissues Network (SMaHT) developed a Multi-Individual Mixed Sample (MIMS) consisting of an *in vitro* mix of six HapMap cell-lines at different concentrations: HG005 at 83.5%, HG02622 at 10%, HG002, HG02257, HG02486 at 2% and HG00438 at 0.5% abundance. This MIMS was then distributed across the network to five Genome Characterization Centers (GCCs): Baylor College of Medicine (BCM), Broad Institute of MIT and Harvard (Broad), New York Genome Center (NYGC), University of Washington and Seattle Children’s Research Institute (UW). and Washington University (WashU). The MIMS was sequenced with both short-read (Illumina NovaSeq X plus) and long-read technologies (Pacific HiFi Revio, and ONT PromethION 24). Bulk RNA-seq were performed with Illumina NovaSeq SP platform. All the data was processed and released by the Data Analysis Center (DAC) at Harvard Medical School. Details regarding library preparation and sequencing are described elsewhere. All the raw and processed files of sequencing data are accessible at https://data.smaht.org/.

### Read mapping and statistics

All data sets were aligned to the GRCh38 reference genome without ALT and decoy sequences. Illumina paired-end reads were mapped with Sentieon’s BWA-MEM (v0.7.17; Sentieon version v202308.01)^41^. Duplicate reads were marked with Picard (v2.9.0)^42^; indel realignment and base-quality score recalibration were performed with Sentieon. PacBio HiFi reads were aligned with pbmm2 (v1.13.0, --preset CCS)^43^, and ONT reads with minimap2 v2.26 (-x map-ont)^44^.

We uniformly sampled 1,000,000 GRCh38 positions (without replacement) to estimate sequencing coverage for Illumina, HiFi and ONT. For each data set, coverage was computed as the number of aligned reads across the sampled positions, from which we derived the depth distribution and the average coverage.

Average molecular length and its distribution were computed per sequencing replicate: for Illumina, from the absolute template length of properly paired primary alignments; for HiFi and ONT, from per-read length.

### Droplet Digital PCR for HapMap Sample Mix Validation

A competitive TaqMan probe approach was used to validate the sample concentrations for the HapMap sample mixture used for benchmarking studies using the method we recently described^45^. In this method, two short fluorescent probes are designed; one for the reference allele and one for the variant allele, each labeled with a fluorescent tag. (FAM and HEX or VIC). A single pair of forward and reverse primers are used for amplification. For each 20□µL reaction, the mix includes 10□µL of ddPCR Supermix for probes (no dUTP), 1□µL of 5□µM forward primer, 1□µL of 5□µM reverse primer, 1□µL of 2.5□µM FAM probe, 1□µL of 2.5□µM HEX probe, an appropriate restriction enzyme (5 units), and 25–100 ng of genomic DNA, topped up with RNase/DNase-free water to the final volume (20-22 µL). The reaction is thoroughly mixed, incubated briefly at room temperature, then partitioned into nanoliter-sized droplets using a droplet generator and transferred to a 96-well PCR plate.

Thermal cycling starts with enzyme activation at 95□°C for 10 minutes, followed by 40 PCR cycles of denaturation at 94□°C for 30 seconds and annealing/extension using a temperature gradient between 50–65□°C for 1 minute to optimize probe and primer binding. A final enzyme deactivation is performed at 98□°C for 10 minutes, then the plate is held at 12□°C until droplet reading. The amplified droplets are read using a droplet reader (BioRad QX200), and the QuantaSoft software analyzes the fluorescence signals from each droplet to classify them as positive or negative for each fluorescent probe signal. Clear separation of positive and negative droplets enables quantification of variant and reference signals. The fractional abundance of the variant is calculated by dividing the number of variant-positive droplets by the total number of droplets positive for either the variant or reference probe, providing a measure of the variant’s VAF level or mixing fraction within the sample.

### Tandem Repeat Allele Delta

Tandem Repeat (TR) Allele Delta is a procedure where for any given TR region in the reference, variants or sequence alignments can be processed to identify changes to a region’s span which typically correspond to TR expansions or contractions. For example, a single 10bp insertion on a haplotype within the region has an allele delta of +10bp. Another example is a read with an alignment that spans the entire region and contains two 5bp deletion cigar operations has an allele delta of −10bp. TR regions were selected from the adotto TR catalog^14^ for being on autosomal chromosomes, having NS annotation of ≥ 1 (i.e., evidence of expansion/contraction), not being interspersed repeat sequence, and regions which were fully spanned by all HPRC haplotype assembly alignments. For the selected TR regions, expected allele deltas were generated for each HapMap sample’s haplotypes using a custom script which parsed the HPRC assembly alignment VCF. Observed allele deltas were generated for each long-read sequencing experiment using qdpi (https://github.com/BCM-HGSC/qdpi). This program parses SAM/BAM/CRAM alignments over a provided BED file of regions and collects total sequencing depth and allele delta per-read. Qdpi was run with the parameter for buffer set to zero, minimum mapQ of reads set to 5, and mapFlag set to 3840. The mapFlag parameter ignores BAM alignments which are not primary alignments (0×100), from reads that fail platform/vendor quality checks (0×200), reads that are PCR or optical duplicate (0×400), and supplementary alignments (0×800).

With allele deltas collected, custom scripts united the expected and observed allele deltas with the following process. For each TR locus, the observed allele deltas were binned to the expected allele delta which was closest by length. For example, a locus with a +15bp and +30bp expected allele delta would collect all observed allele deltas ≤+22 and ≥+23, respectively. Next, expected allele deltas were merged per-locus such that any two allele deltas within 2bp were consolidated. For example, a set of expected allele deltas of +4bp, +6bp, and +8bp would all be consolidated into one by summing their number of observed allele deltas. The number of observed allele delta (i.e., total read support) was divided by the total observed coverage for the entire locus to calculate the observed VAF. Finally, to limit the influence of small indel sequencing errors on results. only regions with alternate alleles ≥5bp were analyzed.

### Modeling VAF, Coverage, and Recall

To model the probability of detecting an allele at a given site, we used a beta-binomial process that accounts for overdispersion in read count data. Specifically, for a given variant allele fraction (VAF) and total read coverage, we computed the probability of observing at least k supporting reads (where k is a user-defined minimum threshold) using the cumulative beta-binomial distribution. The model assumes a beta prior on the binomial probability parameter, with shape parameters chosen to keep the distribution centered on the expected allele fraction while allowing dispersion to vary according to site-specific sequencing depth.

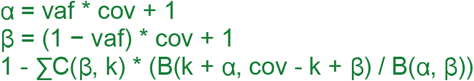

Where vaf is the variant allele fraction and cov the coverage to model, C is the binomial coefficient, and B is the beta function. The summation is performed for each integer i from 0 to (k − 1) to calculate cumulative probability of observing up to k reads. The final probability of observing an allele of the given vaf with at least cov reads is then 1 minus this cumulative value.

To evaluate the fit of each sequencing experiment to the model, we computed the log-likelihood of the observed alternative read count at each site given the total coverage and expected allele fraction. Likelihoods were calculated under both a standard binomial model and the above beta-binomial model, the latter allowing for overdispersion in read counts. Site-specific log-likelihoods were summed across all loci to yield the total log-likelihood for each experiment.

### Benchmark VCF creation

Human Pangenome Reference Consortium assemblies for the six HapMap samples were procured from https://github.com/human-pangenomics/HPP_Year1_Assemblies. Each of the twelve haplotypes were aligned to GRCh38^46^ and variants called with minimap and paftools, respectively^28^. To consolidate variants across haplotypes, BCFtools v1.19^47^ merge was used and custom code consolidated genotype information per-sample. The Variant VCF entries were annotated with SVLEN and SVTYPE info field using Truvari v5.3^31^ anno svinfo, tandem repeat annotations with truvari anno trf (a wrapper around TandemRepeatsFinder^48^ and the adotto TR catalog^14,49^), and annotations of the number of SV neighbors within 1kbp with truvari anno numneigh.

To ensure no information loss while still performing SV merging, truvari phab was used. This program takes as input phased genotypes from regions of a VCF to recreate the original haplotypes described by VCF entries before running a multiple-sequence alignment (MSA) and re-calling variants. We ran truvari phab --buffer 0 on all chunks of the genome with at least one ≥50bp SV as identified by truvari anno chunks. The regions identified were expanded by 100bp and merged with bedtools slop and merge.

With the SVs losslessly harmonized, highly similar, distinct SVs were collapsed with a custom script which reuses the Truvari API and is available in this project’s code repository under analysis/scripts/dynamic_collapse.py. This script analyzes neighborhoods of variants within 1kbp, and collapses SVs with the same type, start position within 100bp, and size similarity above a dynamic threshold. This dynamic thresholding scales the size similarity percent according to the size of the variants being compared based on a linear interpolation formula of:

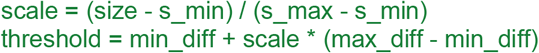

Here, the variant **size** is the smaller of two SVs being compared and the parameters for the dynamic threshold curve were **min_diff**=5, **max_diff**=30, **s_min**=50, **s_max**=1500.

### Benchmark BED creation

To generate the high-confidence bed file, the original alignments generated above were processed and coverage bed files of alignment depth per-haplotype were created. Regions covered with a single alignment on autosomes were extracted. To consolidate singly covered regions, BEDtools v2.31^50^ genomecov was used and reference spans with exactly 12x coverage (1x per-haplotype) were extracted. Reference genome annotation tracks from GRCh38 were downloaded from UCSC Genome Browser^51^. The reference gaps bed file was expanded by 5kbp using the BEDtools slop command. All segmental duplication regions^52^ and simple repeats regions spanning ≥10kbp which were not completely covered by the 12x-bed were extracted using the BEDtools intersect command). All regions with either failed truvari phab variant harmonization (N=3, span=118,378 bp), that produced an increase in SV counts (N=1,898, span=1,927,831 bp), or produced only resized/split variants <50bp (i.e. phab ‘removed’ SVs; N=4,069, span=1,163,929 bp) were also excluded from the benchmark. Next, the original, unmanipulated VCF and the dynamically collapsed VCF were analyzed independently with a custom script to identify regions of high SV density/complexity for each VCF. This process analyzed SVs with size between 50-100kbp using the Truvari API to identify chunks of SVs. A chunk of SVs is defined such that the upstream most SV’s start position and downstream most SV’s end position is at least 1kbp away from any other SV’s start/end while all SVs internal to the chunk have at least one neighboring SV within 1kbp. If any haplotype within a given chunk produced more than one SV, the region was classified as a high density/complexity region. These regions were then expanded by 500bp using the BEDtools slop command. The expanded gaps, uncovered segmental duplications, uncovered large simple repeats, phab impacted, and identified high complexity/density regions were consolidated with BEDtools merge before being subtracted from the 12x-coverage bed to create the final high-confidence bed file. Note that since the last criteria (high complexity/density) is dependent on the VCF being analyzed, two high-confidence bed files corresponding to the two available VCFs were made.

### Droplet Digital PCR (ddPCR) Validation of SV Calls

A subset of SV calls were manually inspected within IGV to generate a breakpoint sequence alignment of a given duplication or deletion. Two TaqMan probes were designed to selectively amplify either the breakpoint junction sequence (FAM) or the wild-type reference sequence (HEX) along with accompanying forward and reverse primers ^53^. The primers and probes were multiplexed into a ddPCR reaction using a standard TaqMan protocol ^54^ on the Bio- Rad QX200 system (Hercules, CA). BioRad ddPCR super mix for probes (No dUTP) was used within the PCR reaction with 25 ng of DNA from the HapMap mixture, and a BioRad C1000 Thermocycler, with the following thermocycling conditions: 95 °C, 58 °C, 72 °C for 35 cycles. The individual FAM and HEX signals from each droplet were then determined with the QuantaSoft Analysis Pro Software to quantify the fractional abundance of the breakpoint sequence within the original sample mixture.

### SV calling

Eight SV callers were tested in this benchmark, as they met two inclusion criteria: (1) the caller supports single-sample input (no matched control required) and (2) supports VAF <10% detection under default settings. For Illumina, we ran Delly (v1.2.6)^33^, SvABA (v1.2.0)^38^, and GRIDSS2 (v2.13.2)^34^. For long reads, we tested Delly (v1.2.6), Sniffles (v2.4)^11^, Severus (v0.1.1)^37^; HiFi-specific softwares included SVDSS (v2.0.0-alpha.1)^39^, pbsv (v2.9.0)^35^, and Sawfish (v0.12.4)^36^. All callers were executed with default parameters unless noted. For Sniffles, we merged outputs from –germline and –mosaic modes and set –mosaic-af-min 0.001 and –mosaic-af-max 0.22 following the developer’s suggestion. VCF normalization was carried out for each call set to produce one SV per record, only FILTER=PASS entries retained, and evaluation restricted to deletions, insertions, and duplications that are ≥50 bp.

### Performance evaluation of somatic SV calling

Each call set was benchmarked against the MIMS benchmark set with Truvari using default parameters and –includebed set to the high-confidence regions. Truvari matches baseline and comparison sets of SVs of the same type between 50bp to 50kbp using thresholds of maximum reference distance at 500 bp, and 70% sequence and size similarity. The parameter –dup-to-ins was used to classify duplications as insertions for consistency with the benchmark schema. Importantly, the –pick parameter, which controls the number of matches a call participates in, was left to the default ‘single’ behavior. This ensured each variant was only counted once, which is especially important given the highly similar SVs this mosaic benchmark represents. Baseline variants were considered germline if present in HG005 or somatic otherwise. As most callers do not separate somatic from germline SVs in single-sample mode, we designated caller-reported SVs with the reported VAF<25% as somatic. This VAF threshold safely exceeds the maximum designed pseudo-somatic VAF (16.5%) of the benchmark set. Somatic performance metrics therefore quantify detection of low-VAF events while ignoring germline calls:

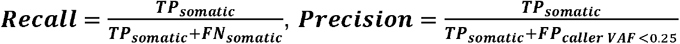

Somatic SVs from the benchmark were partitioned into six VAF bins: <0.5%, 0.5-1%, 1- 4%, 4-5%, 5-10%, and 10-20%, chosen to roughly balance SV counts per bin. VAF-bin-specific recall is calculated based on expected VAF (exp. VAF), and VAF-bin-specific precision based on each caller’s reported VAF (caller VAF): for VAF bin A,

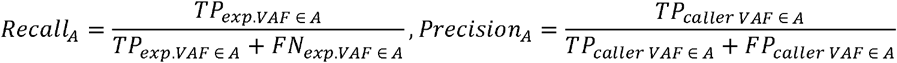

This keeps the VAF calculation consistent within each metric. Note that *TP*_*exp*.*VAF ∈A*_ and *TP*_*caller VAF ∈ A*_ are distinct, and a caller may have an undefined precision for certain bin *A* if it reports no calls with *caller VAF ∈A*.

In the difficulty-stratified analysis, each benchmark SV and each caller-reported SV was annotated with five features: insertions; size ≤250 bp (small SV); low VAF (<10%); breakpoint overlapping a tandem repeat (TR); and proximity to another SV breakpoint within 500 bp (adjacent SV). Variants lacking any of these were assigned to “not-challenging”. Because features can co-occur, difficulty-specific recall was computed on difficulty-exclusive sets (see **Supplementary Figure S8** for comparison with non-exclusive sets). Difficulty-specific precision was computed on the full difficulty sets by subsetting each caller’s calls to the category.

### Platform-specific detection limit evaluation

To compare platforms at matched depth, replicate BAMs were merged per platform using SAMtools (v1.17; samtools merge)^55^, then randomly down-sampled to 60X, 90X, 120X, 150X, and 180X using SAMtools (samtools view -s) for short reads and an in-house sampler for long reads. Coverage after downsampling was verified with SAMtools (samtools depth). The same callers were run on each downsampled data set, and performance was evaluated with VAF-bin-specific recall and precision.

To contextualize caller recall by the proportion of SVs actually detectable in each data set, we estimated an upper bound via platform-specific genotyper on every downsampled BAM. For Illumina, we ran Paragraph (v2.4a)^56^ against the benchmark SV set and each BAM, extracting per-SV supporting-read counts from the genotyped VCF’s FORMAT:AD field. For long reads, we used Sniffles (v2.4)^11^ in force-calling mode with the same inputs and extracted supporting-read count from FORMAT:DV field for each SV. For each data set and support level (≥1, ≥2, ≥3 reads), the upper bound is the proportion of benchmark SVs with at least that many supporting reads.

### StratP Test

To identify variant stratifications that significantly impact a tool’s performance, we developed the StratP test to evaluate whether a caller’s accuracy (e.g., recall) is dependent on a stratification. StratP computes a directional enrichment score using expected counts under independence, and evaluates significance via permutation-based resampling. This is necessary because variant features can co-occur non-randomly. For example, tandem repeats may more frequently contain insertions than deletions, creating imbalanced subgroups that could bias naive stratification analyses. By evaluating enrichment across groups defined by combinations of feature values and assessing significance through permutation, StratP helps mitigate such confounding effects without requiring explicit modeling of feature dependencies. The resulting p-values provide a robust, interpretable measure for prioritizing stratifications that are impactful to a tool.

The StratP test evaluates whether specific variant features are significantly associated with differences in performance (e.g., enrichment or depletion of true positives). It operates on a labeled set of variants (e.g., classified as true positive or false negative by Truvari) along with a set of categorical annotations (e.g., SV Type and Size Bin). Each SV is assigned to a group defined by the combination of its features’ values. For example, groups might correspond to all deletions within tandem repeats, or all insertions outside tandem repeats. The number of true and false SVs in each group is tallied to build a contingency table, and groups with fewer than a minimum number of observations (default: 10) are excluded. Next, the expected counts for each cell in the contingency table are calculated under the assumption of independence between rows and columns. Let:

R_i_ be the total count of row i (i.e., number of false/true variants)

C_j_ be the total count of column j (i.e., number of variants in a group)

N be the total number of observations (i.e., variants)

Then the expected count for cell (i, j) is given by:

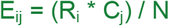

A directional enrichment score is then computed for each group as:

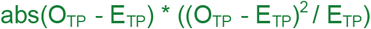

Where O_TP_ and E_TP_ are the observed and expected true positive counts for the group, and abs is the absolute value operator. This score is positive when the group is enriched for true positives, negative when depleted, and zero when observed matches expected. All groups are then ranked according to their enrichment scores.

To test whether a particular feature/value pair (i.e. stratification) is associated with higher or lower enrichment, the difference in mean rank is computed as:

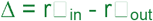

Where r□_in_ is the average rank of groups containing the stratification, and r□_out_ is the average rank of groups not containing it. A permutation test (default: 10,000 permutations) is performed to assess the statistical significance of Δ by shuffling ranks between r_in_ and r_out_ and a new delta collected at each step. A p-value is assigned by comparing the observed Δ to the distribution of permuted deltas according to the chosen test direction:

- One-tailed (left): proportion of permutations where Δperm ≤ Δobs
- One-tailed (right): proportion of permutations where Δperm ≥ Δobs
- Two-tailed: proportion of permutations where abs(Δperm) ≥ abs(Δobs)

Finally, all resulting p-values are corrected for multiple hypothesis testing using the Benjamini-Hochberg procedure to control the false discovery rate. The StratP test is available as a sub-command of Truvari that operates on Truvari bench output directories. Parameters for stratifications from the SMaHT MIMS SV benchmark can be specified with --preset mims. Alternatively, the script exposes parameters to allow testing of custom stratifications and other benchmarks. All benchmark stratification annotations, with the exception of expected VAF, can be placed into VCFs using the truvari anno commands of numneigh and trf.

### Evaluation of orthogonal sequencing

A whole-genome sequencing experiment was performed at the BCM GCC using the Element Bioscience short-read Aviti platform. Alignment was performed using bwa mem (v0.7.15). SV calling was done replicating the analysis pipeline utilized for Illumina short-read experiments with Delly (v1.2.6), SvABA (v1.2.0), and GRIDSS (2.13.2). Genotyping of DEL was performed with SVTyper (v0.7.1), using default parameters.

Bulk RNA-seq experiments were performed by the four GCCs using Illumina short reads with a target of 150 million reads. Details regarding library preparation and sequencing are described elsewhere. All sequencing data was analyzed by the DAC which reported transcripts per-million (TPM) and Fragments Per Kilobase-Million (FPKM) by gene and isoform/transcript.

For each replicate, we used all genes which had TPM or FPKM values greater than zero (expressed genes). We then overlapped the exons of the expressed genes with the SVs of the benchmark, and classified the SV as candidate for validation if It covered at least 50% of any exon with an SV (DEL) and was within the gene body if the SV is an insertion (INS). All candidate SVs were manually curated in IGV

## Data availability

All data are openly available at phs004193 and at https://data.smaht.org/.

## Code availability

All scripts and the benchmark can be found at https://github.com/BCM-HGSC/SMaHT_MIMS/.

## Acknowledgement

Research reported in this publication was supported by the Office of The Director, National Institutes of Health under Award Numbers UM1DA058230, UM1DA058229, UG3NS132105 and UM1DA058220.

## Author contribution

F.J.S. and P.J.P. supervised the project. A.C.E. generated the reference SV set and regions and improved computational methodologies. Y.Z. performed variant calling, benchmarking, and detection-limit analyses. L.F.P. carried out orthogonal sequencing-based validation. C.M.G. performed orthogonal validation using ddPCR. S.M. contributed to the design and implementation of StratP. Y.Z., A.C.E., F.J.S., and L.F.P. drafted the manuscript. Y.Z., A.C.E., L.F.P., T.M., P.J.P., and F.J.S. revised the manuscript with input from all authors. All authors read and approved the final version.

