## Supplementary Text for "Comprehensive benchmarking of somatic structural variant detection at ultra-low allele fractions"

### Section 1

#### HPRC Assembly Details

The per-haplotype assemblies contained an average of 379 contigs (standard deviation ±145) averaging 3,014,355,355 bp (±44,788,834) in length. The average contig N50 was 61,643,268 (±11,458,838) and L50 of 17 (±2).

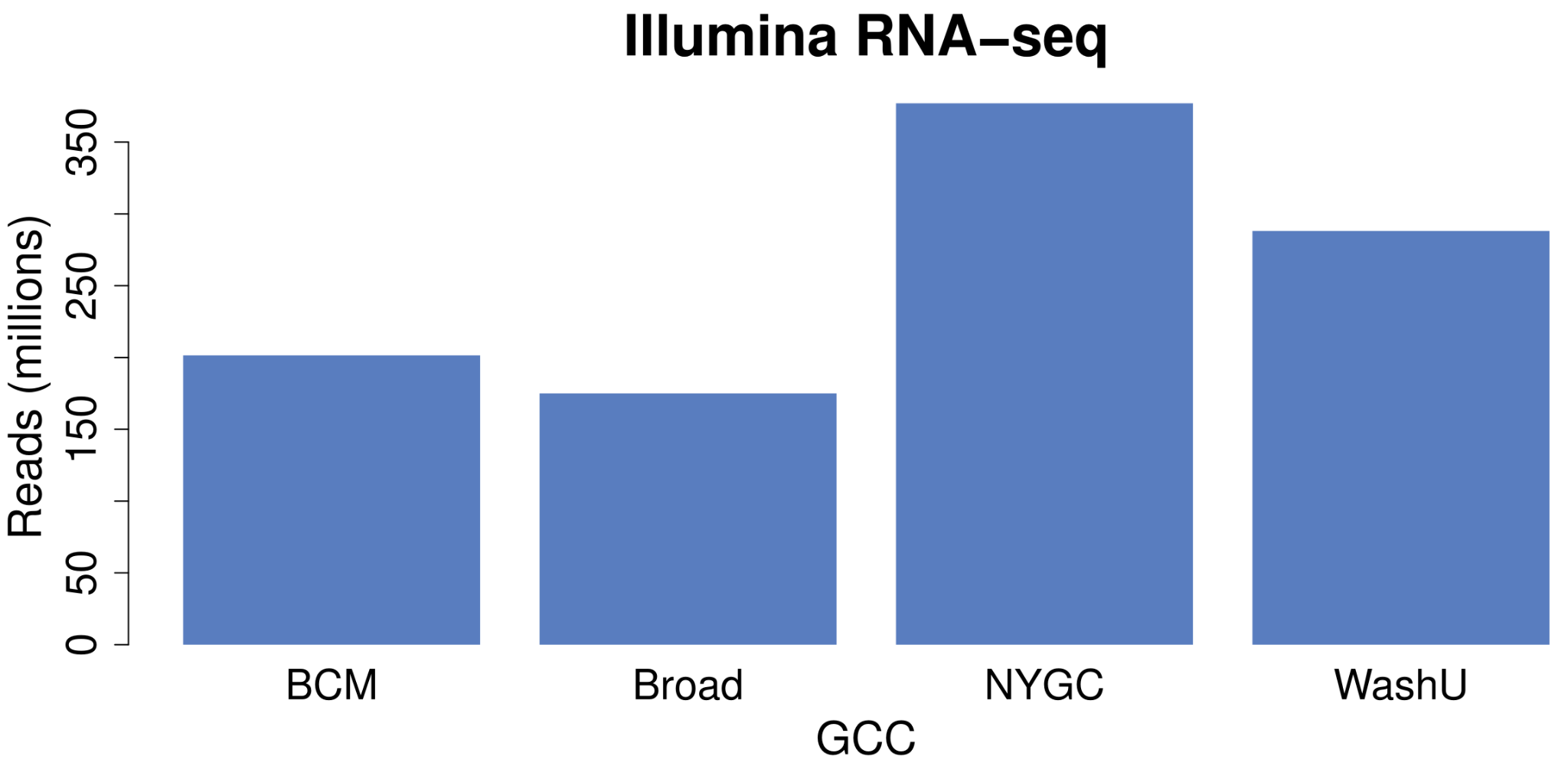
***Supplementary Figure S1*** *- Sequencing statistics of the bulk RNA-sequencing experiment by Genome Characterization Center (GCC). Target sequencing depth is 150 million reads, with all GCCs submitting higher sequencing depth.*

#### TR Allele Delta

Allele Delta is a technique where sequences which align over tandem repeats are analyzed and expansions/contractions are identified. For example, a read which aligns over a tandem repeat and gives rise to a 10 bp and 12 bp insertion has an allele delta of +22 bp. Allele deltas from the HapMap assembly haplotype were collected as the benchmark allele delta. Next, for each locus, the benchmark allele deltas were consolidated when the difference in their deltas was less than 2 bp. The process of consolidation included summing expected VAFs of each haplotype’s alleles based on the HapMap mix proportions. Only benchmark allele deltas greater than 5 bp were analyzed. Each available long-read sequencing experiment was processed with qdpi (<https://github.com/BCM-HGSC/qdpi>) to collect each read’s allele delta per-TR locus. Only primary alignments with a minimum MAPQ of 5 were considered. Next, each read allele delta at a locus was assigned to the benchmark allele delta that was closest in length. The number of reads assigned to each benchmark allele delta divided by the total coverage of reads spanning the TR locus created the observed VAF. Only benchmark allele deltas with at least one supporting read were analyzed.

A total of 573,057 tandem repeats from adotto were analyzed based on criteria of being within the SMaHT MIMS benchmark regions as well as having adotto annotations of >1 sample and non-interspersed repeats. After filtering to only loci with at least 1 HapMap sample having a ≥5bp expansion/contraction, 48,910 TR loci remained for analysis. For each sequencing experiment, loci with no reported coverage of reads spanning the region or without all alternate alleles having ≥5bp expansions/contractions were excluded to limit noise from small variants and/or small sequencing errors. The coverage distribution across TR alleles in **Supplementary Figure S2a**. The difference in observed and expected VAF for each sequencing experiment is in **Supplementary Figure S2b**. A Bland-Altman plot (**Supplementary Figure S2c**) was created to investigate any VAF measurement bias. Overall, nearly zero bias was observed. However positive bias (increased observed VAF) in low-VAF alleles and negative bias (decreased observed VAF) in high-VAF alleles was evident. These directional biases with respect are a natural consequence of the sampling floor. For example, given 100x coverage, an allele with an expected VAF of 0.025% can be observed at a minimum of 1% VAF. Furthermore, the somatic sampling floor combined with competition for sequencing coverage of alleles causes alleles in the high VAF (i.e. germline homozygous alternate) to be undersampled.

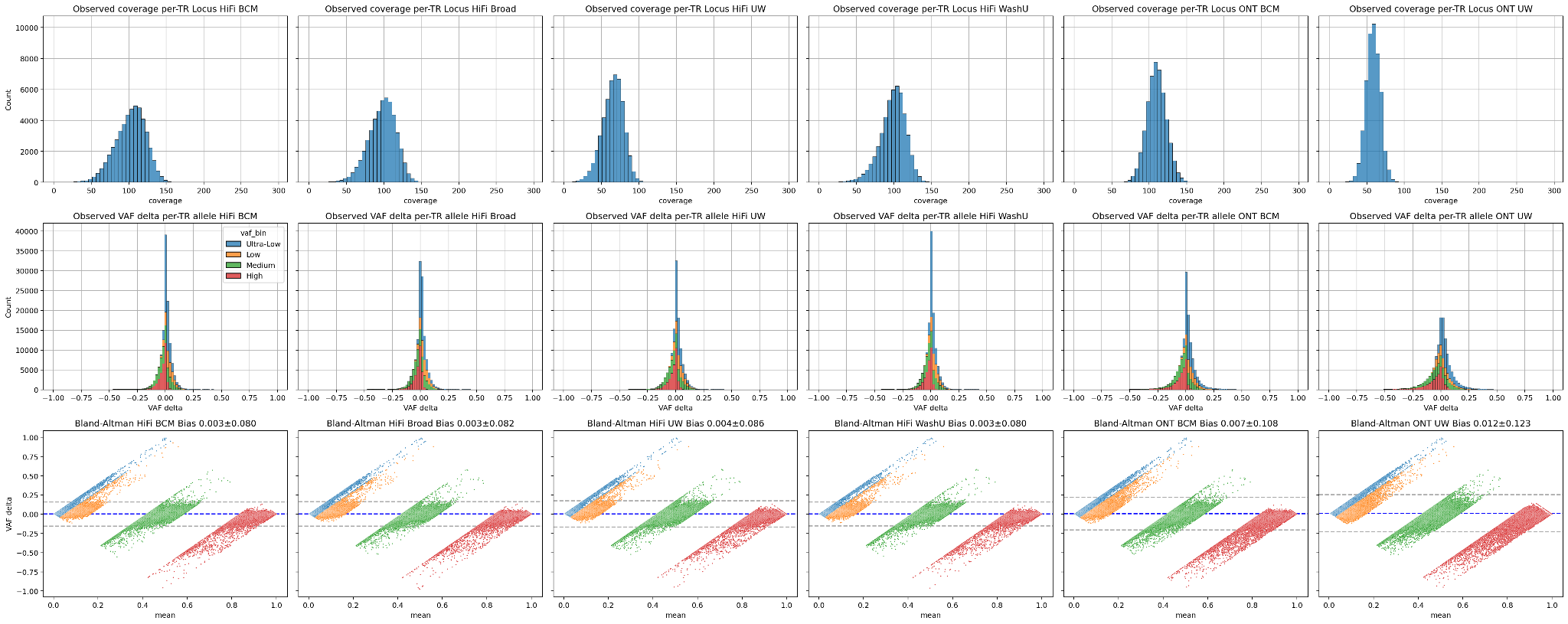

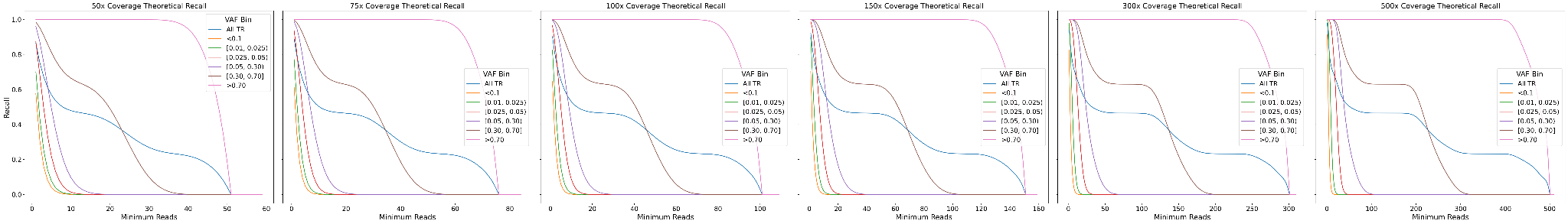

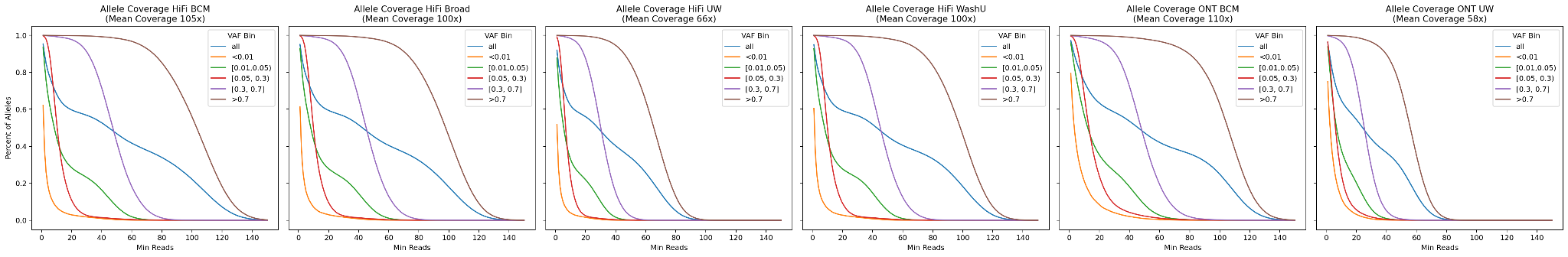

a)

b)

c)

d)

e)

***Supplementary Figure S2:*** *- Allele Delta details a) Average coverage per-TR locus across long-read sequencing technology and sequencing center. b) Distribution of expected minus observed VAF for each TR allele across centers and technologies. c) Bland-Altman plots of observed and expected VAF mean (X-axis) against the observed-expected VAF difference (Y-axis) Hues correspond to VAF bins. d) Empirical cumulative distribution functions of sequencing experiments’ TR recall as a function of minimum number of read support overall and by VAF bin. e) Cumulative Distribution Functions of Beta-binomial process modeling recall as a function of min support thresholds for (left to right) 50x, 75x, 100x, 150x, 300x, 500x coverage.*

### Section 2

#### SV Merging

As described in the main results, discovery of SVs from the HPRC assemblies involved three major steps: Initial alignment and variant calling from minimap2 and paftools; harmonization of SVs using truvari phab; collapsing of highly similar SV alleles. The first transformation of SVs from the original, independent alignments to harmonized multiple-sequence alignment resulted in a reduction from 54,369 SVs to 47,638 SVs (**Supplementary Figure S3a**). Because redundant SVs can be placed at different positions, we also observed that the number of positions at which an SV occurs drops from 36,102 in the original VCF to 28,904 in the harmonized VCF. These reductions in SV count and SV positions are the result of a more parsimonious description of the set of SVs. To illustrate the impact of SV harmonization versus classic SV merging, we also ran truvari collapse on the original VCF and measured the SV count at 39,077 and SV position count at 33,443. Truvari collapse produced fewer SVs than phab harmonization, however the SV positions were greater and nearer the number of SV positions seen in the original alignments (**Supplementary Figure S3b**). This is due to truvari collapse picking representative SVs for sets of matched variants which are spread out across multiple positions in the original alignment space, but those same SVs become placed at the same position with phab. For the final benchmark, the phab harmonized SVs were still collapsed using a dynamic collapsing procedure. This produced fewer SVs, however the number of SV positions was consistent to those seen in the phab harmonized SVs. The cause of fewer SVs from phab+collapse compared to just truvari collapse is largely due to looser similarity thresholds for smaller SVs. Dynamic collapsing capped the size difference between two SVs at between 5bp to 30bp whereas truvari collapse used a flat 95% similarity threshold. For SVs less than 100bp, 95% is more stringent than the 5bp dynamic collapse. For example, 50bp SVs will dynamic collapse with SVs ±5bp, whereas truvari collapse will match SVs ±2bp.

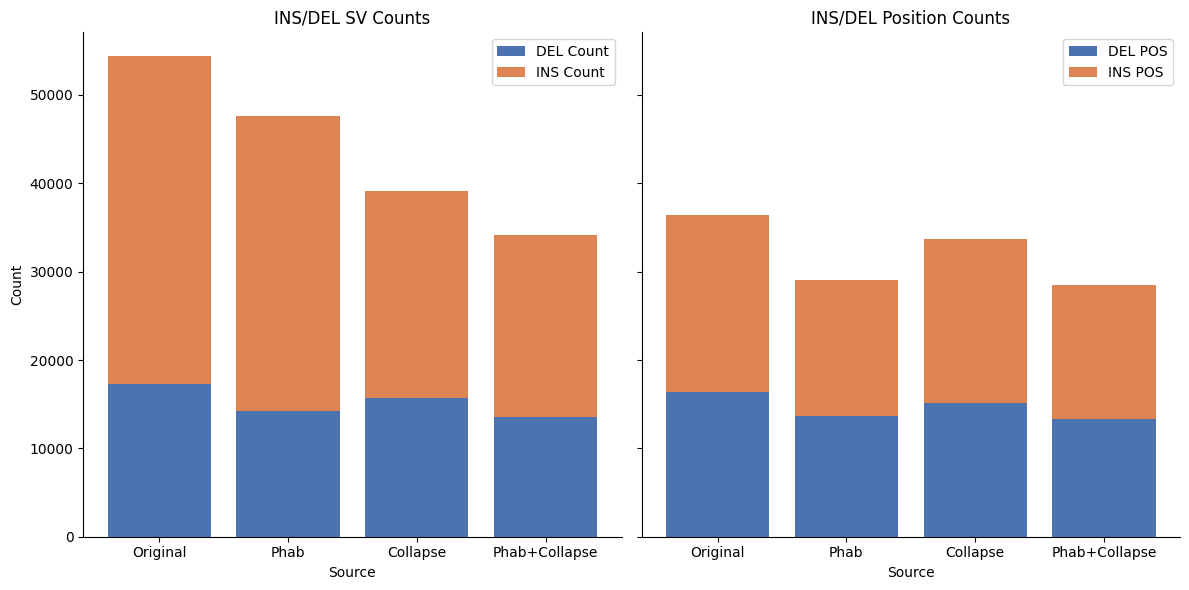

a)

b)

***Supplementary Figure S3 -*** *a) SV counts for three merged sets of SVs - original from bcftools, phab harmonized, truvari collapse, phab harmonized and dynamic collapsing. b) SV position counts for three merged sets of SVs - original from bcftools, phab harmonized, truvari collapse, dynamic collapsing.*

While SV collapsing procedures are inherently lossy they do have situational advantages. The largest advantage is preservation of SV representations. Phab harmonization produces SV representations from the multiple-sequence alignment that, while certainly more parsimonious, may not reflect the same SV representations that are produced by independent sequence alignment. In practice, alignment of sequencing reads is performed independently per-read. Therefore, the SV representations that reads produce are more consistent to independent alignment of haplotypes than what’s produced by the MSA. In contrast, while SV collapsing may over-merge SVs, the kept representation is guaranteed to be reflective of the original alignment space as the SV caller results that produced it. Furthermore, SV collapsing does not require phased genotypes or the presence of smaller variants. Phab harmonization, however, requires phased genotypes of SVs, and it is highly recommended that all variants which describe a haplotype (i.e. <50bp events) are present to ensure the optimal SV representations are produced. Without these SNPs and small insertion/deletions, the resultant SV representations may not accurately reflect the biological sequence from which they were derived.

**Supplementary Figure S4** - SV Neighbor Count Distribution of benchmark SVs

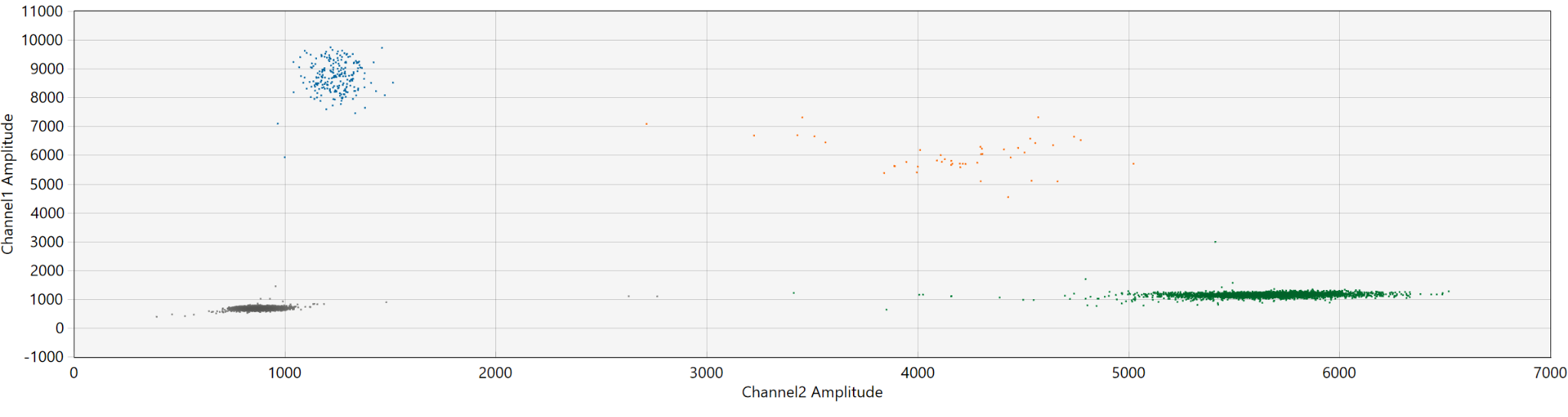

chr3:133357512

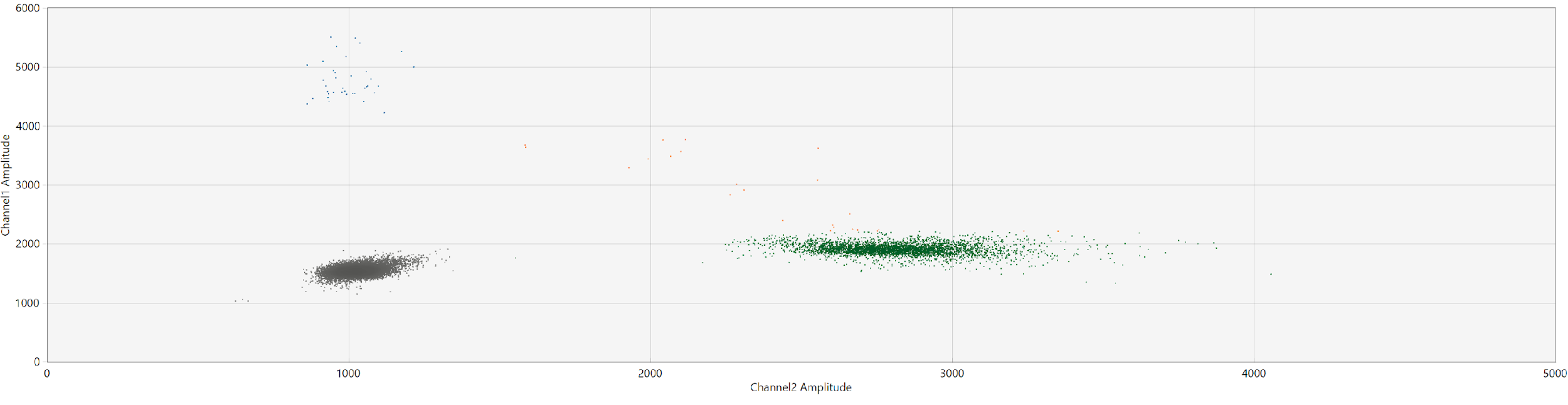

chr20:19189417

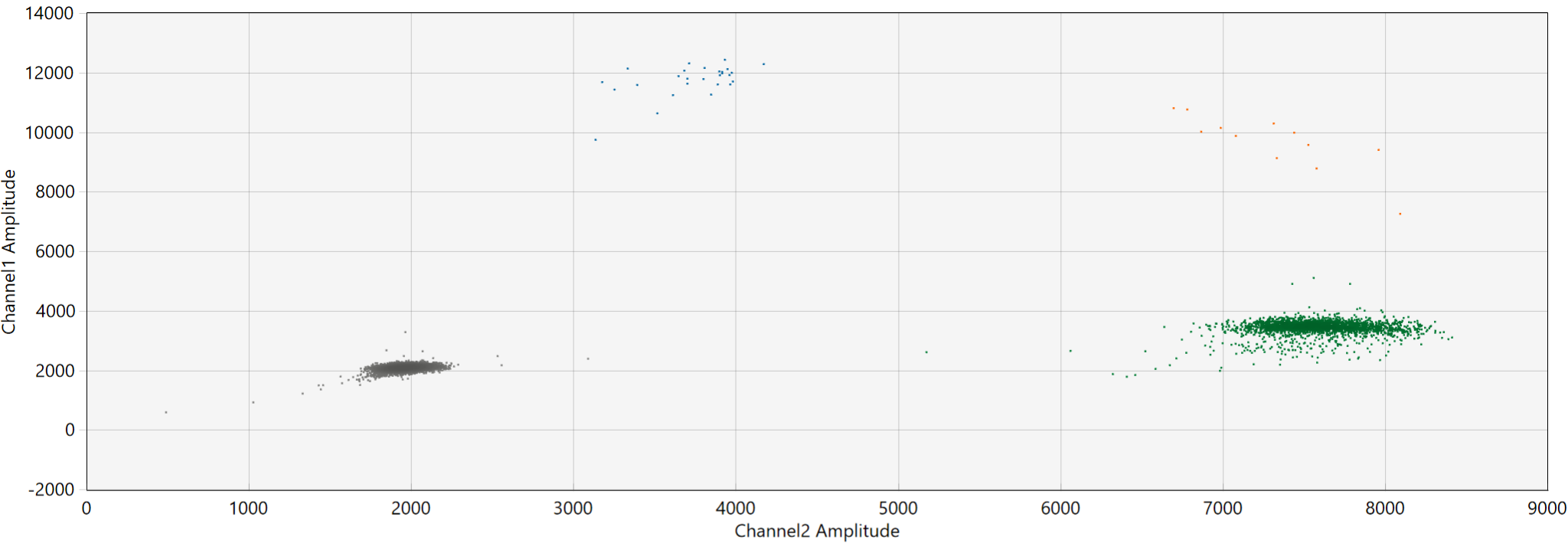

chr12:7680230

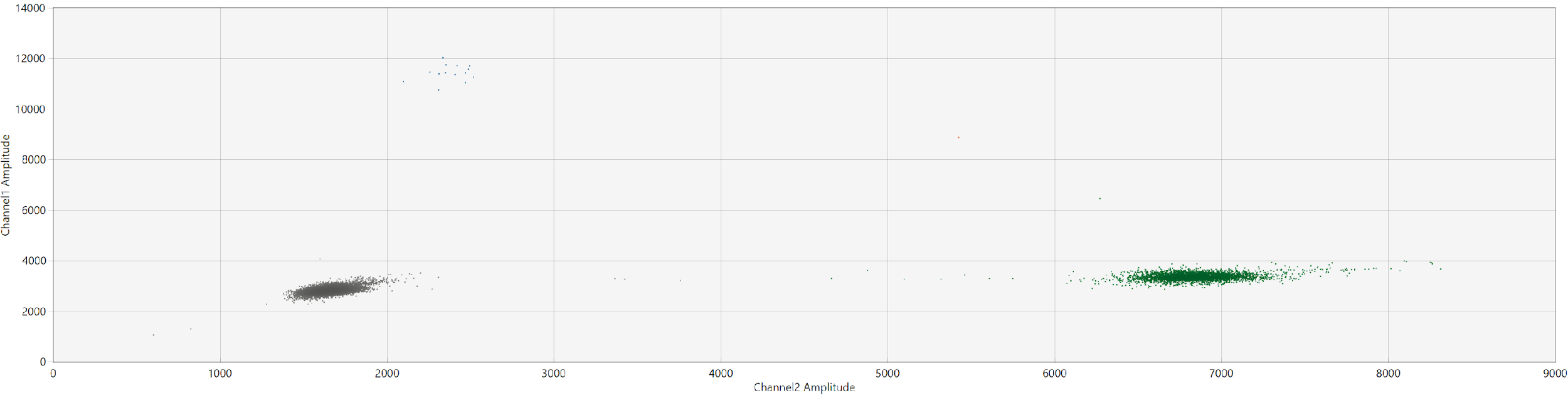

chr2:81703819

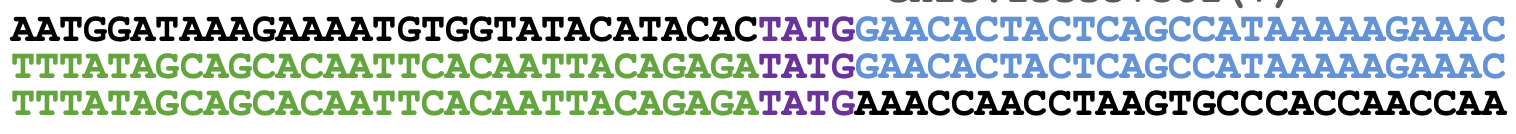

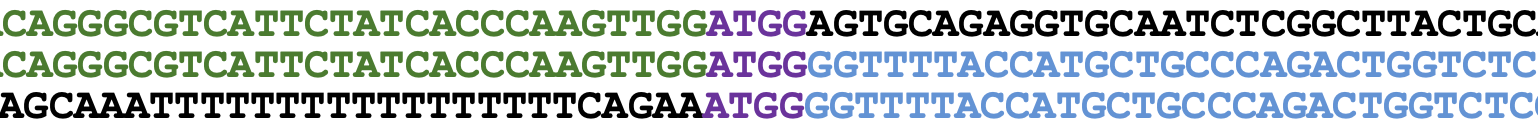

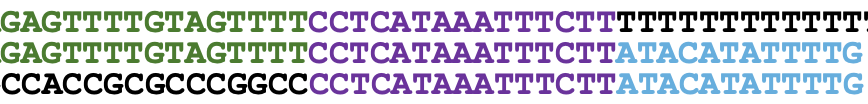

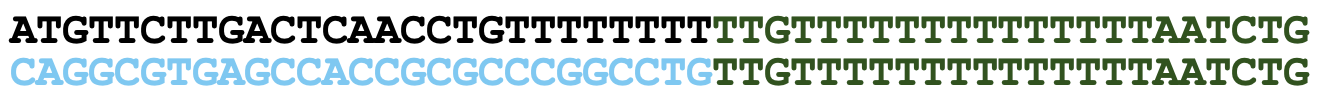

**Supplementary Figure S5 -** digital droplet PCR validation over 4 SVs. Supplementary Table 3 holds details of SVs including size and observed VAF. Above each ddPCR plot is the PCR sequences. Chr2 (d) only shows one side of the breakpoint for the insertion. Scatterplots show channel 2 (x-axis) and channel 1 (y-axis) amplitudes. Black circles highlight the light-blue points of FAM amplified sequences (i.e., SV breakpoint), green points are HEX sequences (i.e., reference sequences). Gray points are negative droplets lacking fluorescence.

### Section 3

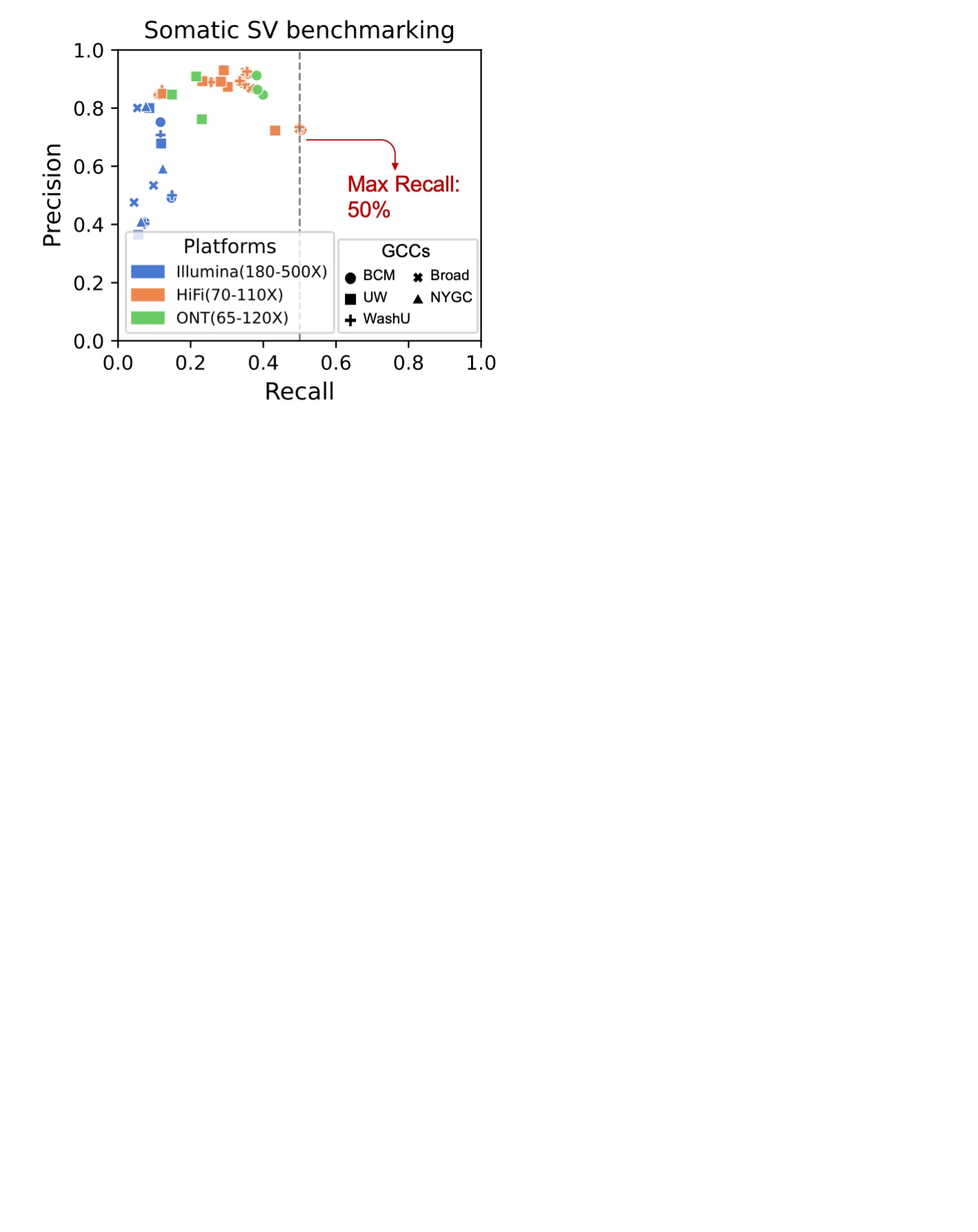

**Supplementary Figure S6** - **Evaluation of 45 SV call sets from 12 somatic SV calling strategies across sequencing replicates.** Recall (x-axis) and precision (y-axis) are shown. Color represents platform (Illumina/HiFi/ONT); marker shape represents GCC replicate; caller identities are not shown.

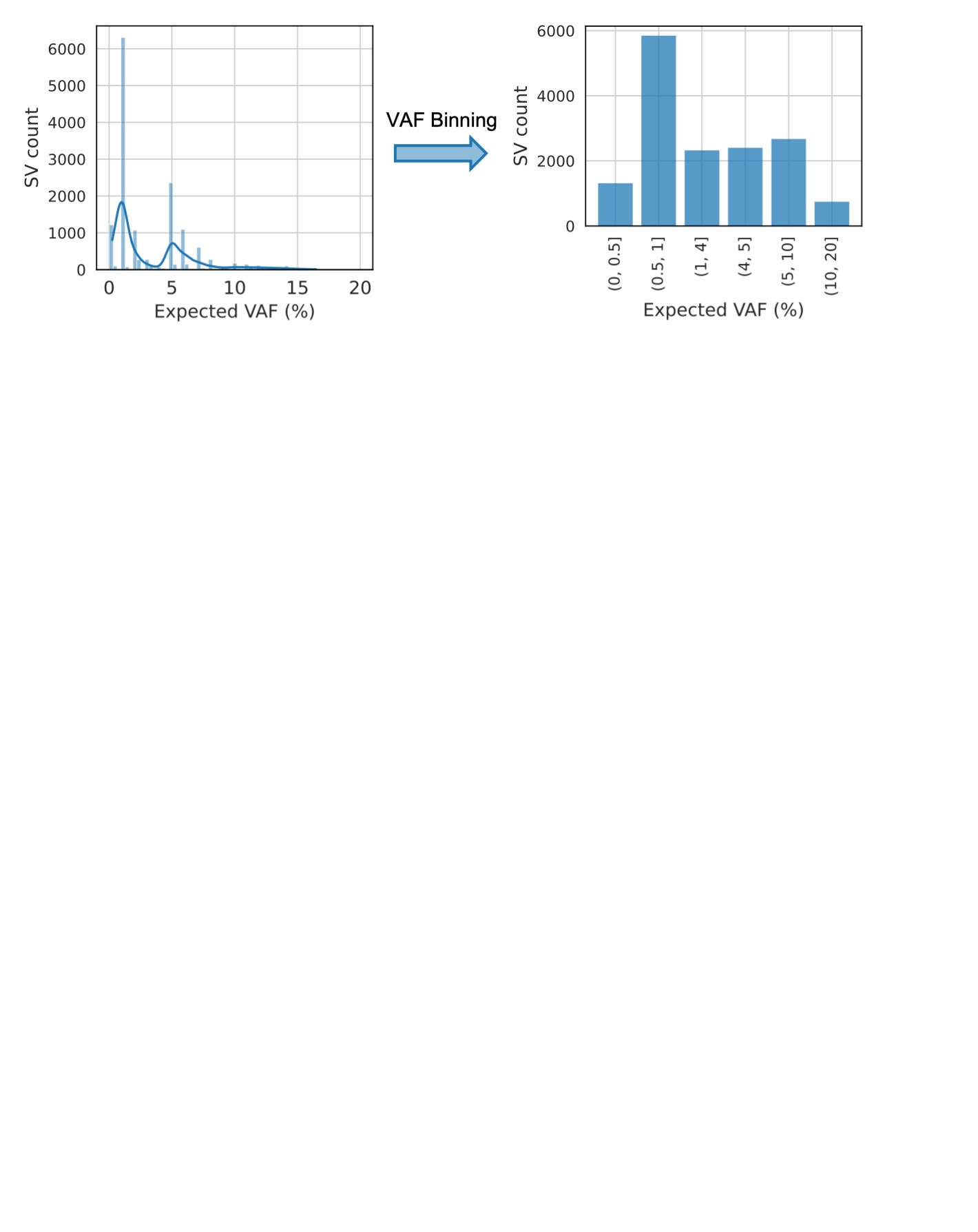

**Supplementary Figure S7** - **Discretization of benchmark somatic SVs by expected VAF.** Left: histogram with 0.05% bin width. Right: counts after grouping into six VAF bins (<0.5%, 0.5–1%, 1–4%, 4–5%, 5–10%, 10–20%) chosen to approximately balance bin sizes.

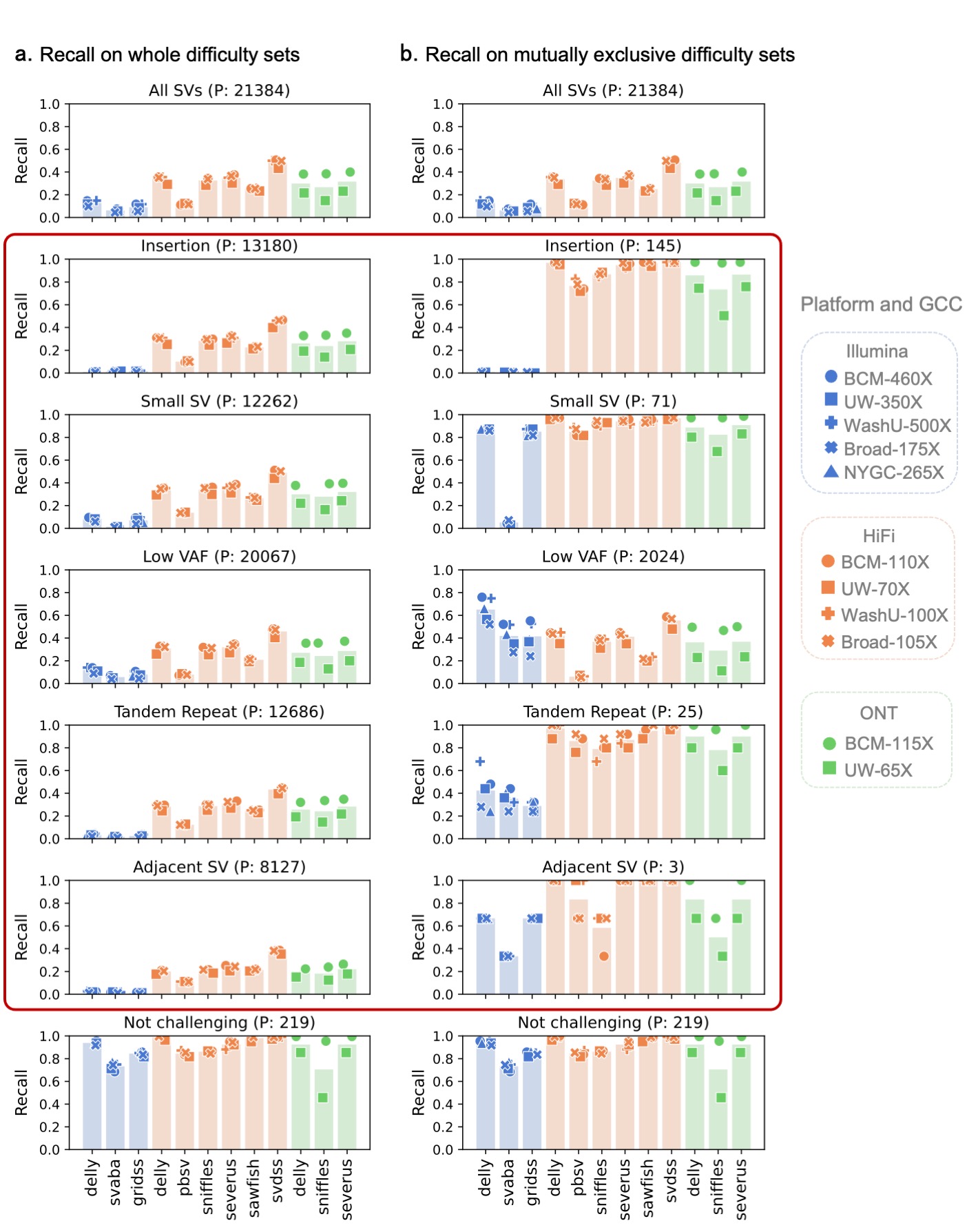

**Supplementary Figure S8** **- Comparison of recall on difficulty vs. difficulty-exclusive sets.** Recall for 12 SV calling strategies across replicates on each **a.** difficulty set (not-exclusive) and **b.** mutually exclusive difficulty set are shown (red block highlights the difference between **a** and **b**). The number of true SVs per set is shown in the panel titles. Points show individual sequencing replicates (color = platform; marker shape = GCC replicate); vertical bars show the replicate mean per workflow. Panel of each difficulty set in **a** resembles the “All SVs” results due to the difficulty co-occurrence; the exclusive stratification reveals platform- and caller-specific advantages.

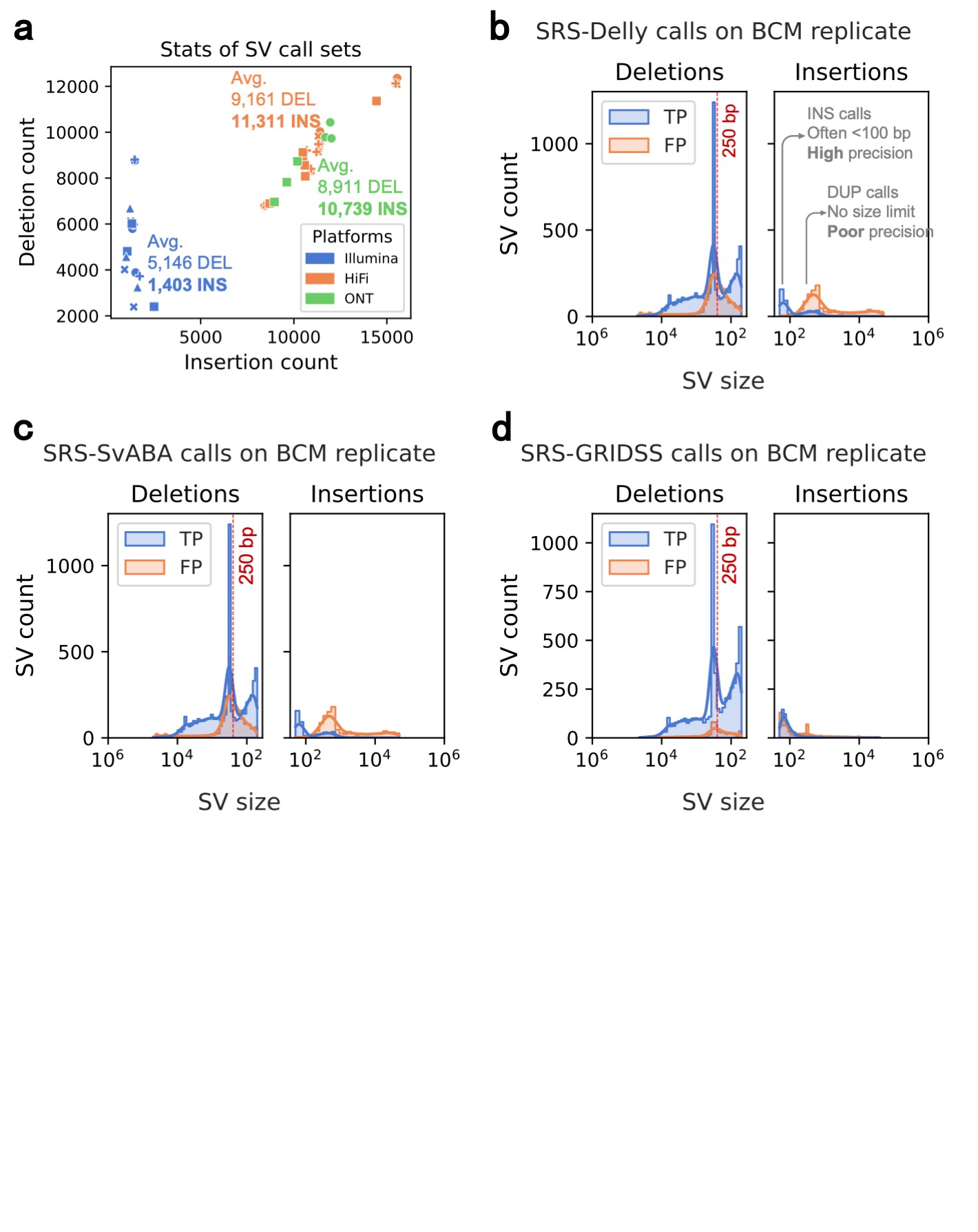

**Supplementary Figure S9** **- SV type and size distributions in short-read (SRS) call sets. a.** Counts of insertions (x-axis) versus deletions (y-axis) for 45 SV call sets, colored by platform; platform-specific mean counts are annotated. b-d. Size distribution of deletions and insertions for three short-read-based callers–Delly (b), SvABA ©, and GRIDSS (d) – evaluated on the same SRS replicate (BCM replicate). True positives (TP) and false positives (FP) are shown in blue and orange, respectively. SV size (bp) is plotted on a log10 scale.

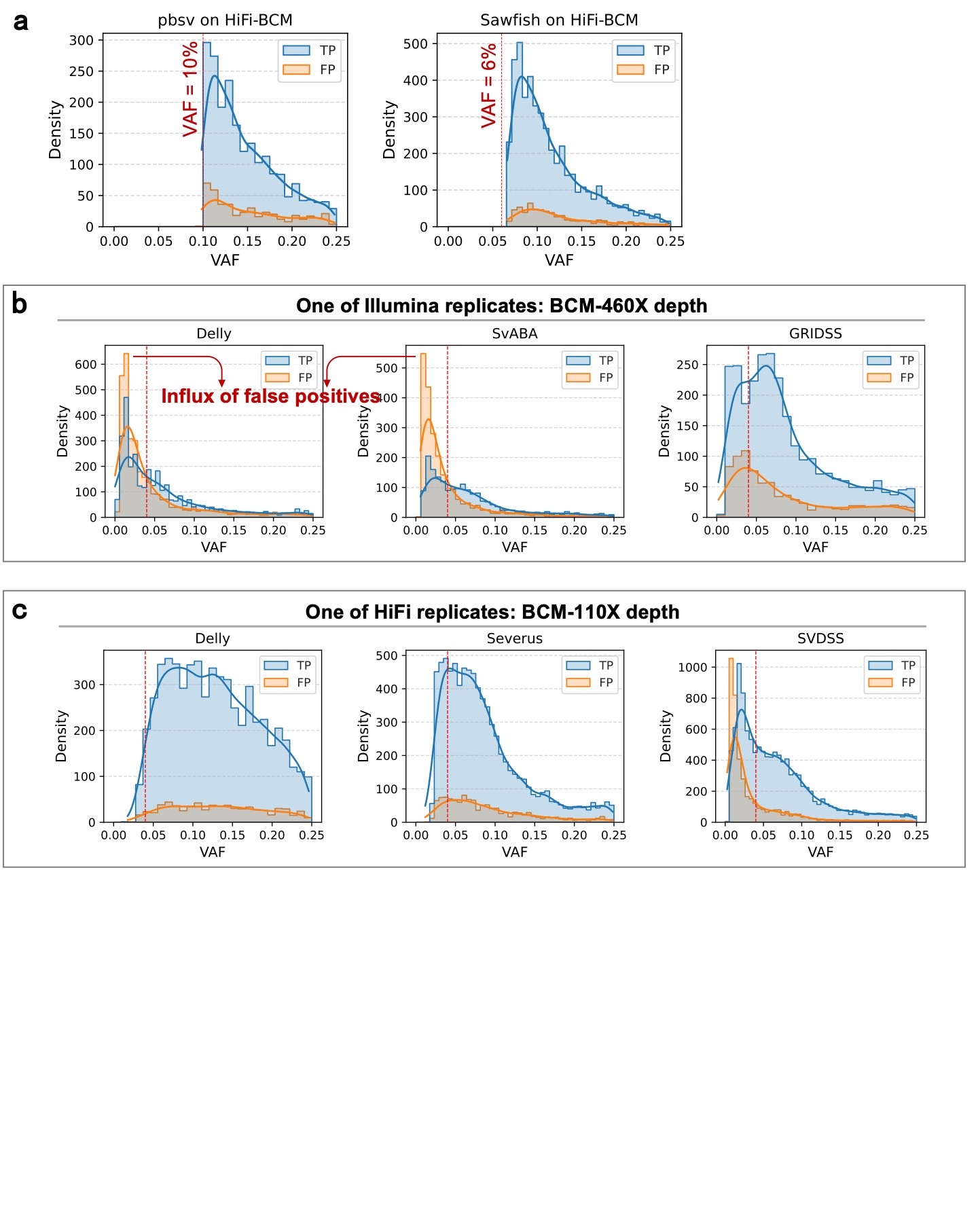

**Supplementary Figure S10** **- VAF distributions of SV call sets, stratified by true positives (TP) and false positives (FP). a.** pbsv and Sawfish on the HiFi BCM replicate: pbsv calls are enriched at VAF > 10 %, and Sawfish at VAF > 6 %. **b.** Three Illumina-based callers on the BCM replicate (460X). **c.** Three HiFi-based callers on the BCM replicate (110X). A red dotted line marks VAF = 0.04 (4 %) in panels **b** and **c**.

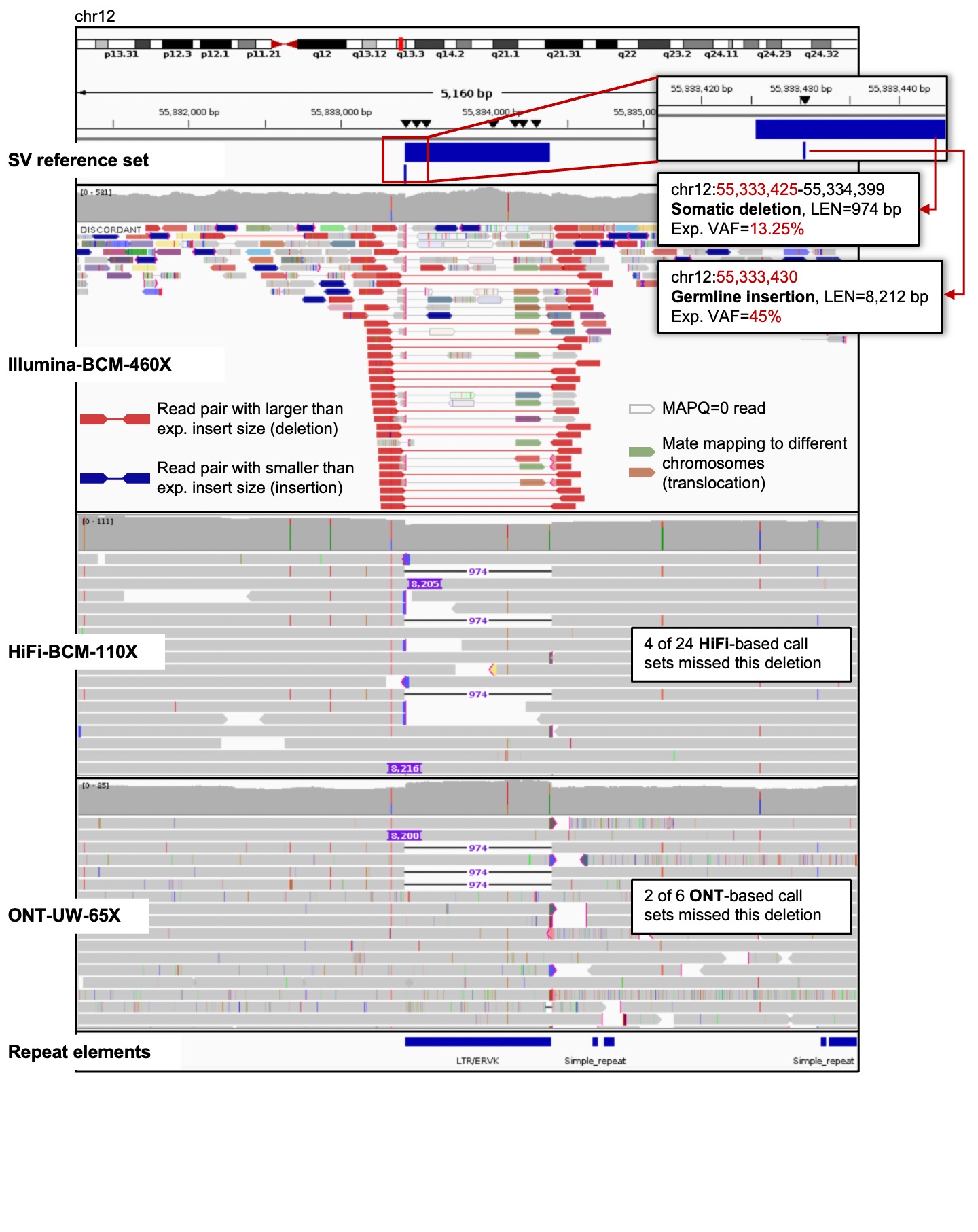

**Supplementary Figure S11** **- IGV snapshot of an adjacent SV across Illumina, HiFi, and ONT alignments.** The left breakpoint of a chr12 somatic deletion lies close to the breakpoint of a germline insertion. This proximity obscures the detection of this somatic deletion.

### Section 4

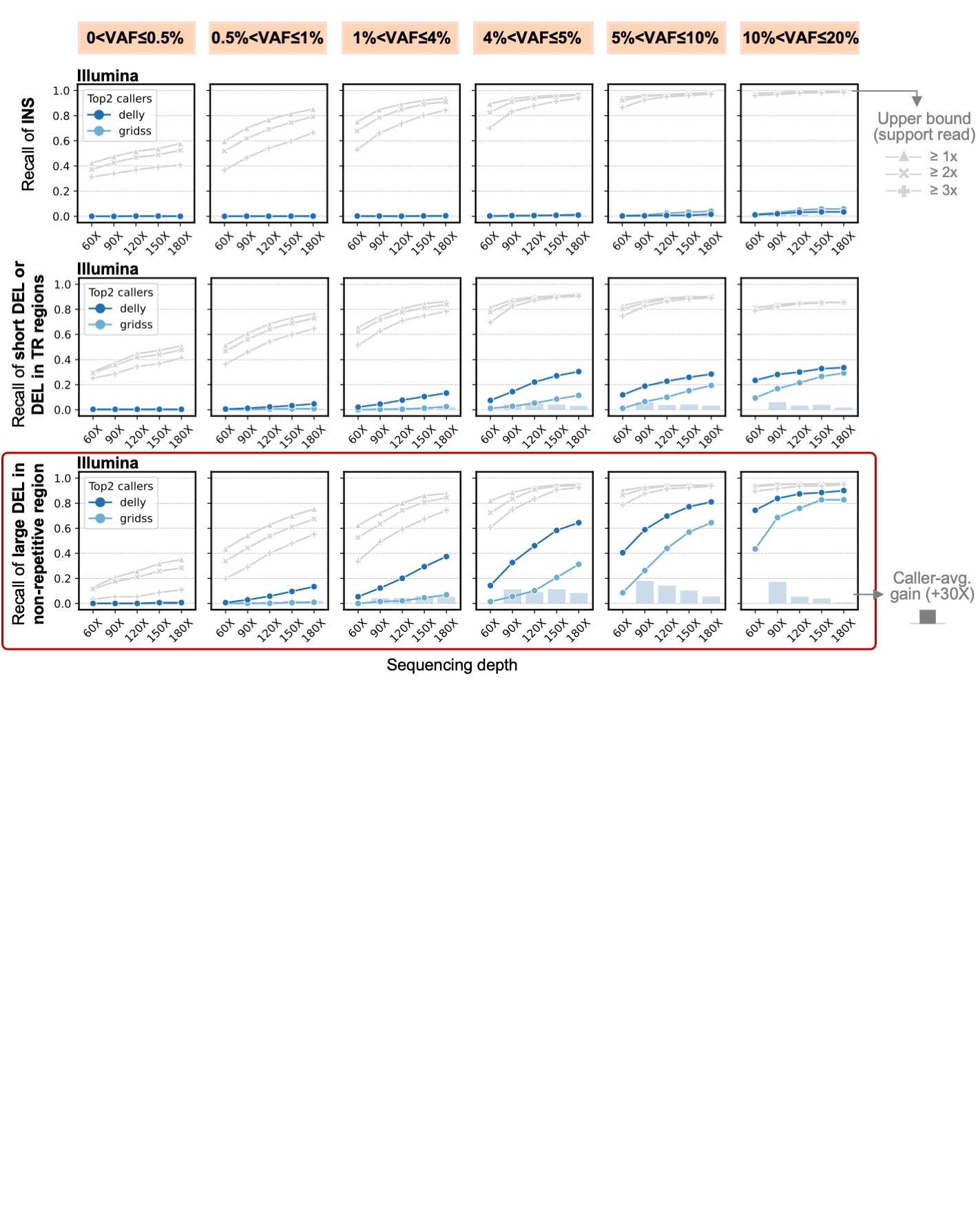

**Supplementary Figure S12 - Illumina-based recall across VAF bins by SV class.** Top: insertions. Middle: deletions ≤250bp or overlapping tandem repeats. Bottom: large deletions (>250bp) in non-repetitive regions. The two best callers (Delly and GRIDSS) are shown; each point is recall in a VAF bin at a given depth. Gray lines show the genotyper-estimated upper bounds requiring 1, 2, or 3 supporting reads. Bars show the average recall gain (across the two callers) for an additional 30X depth increase relative to the preceding depth.

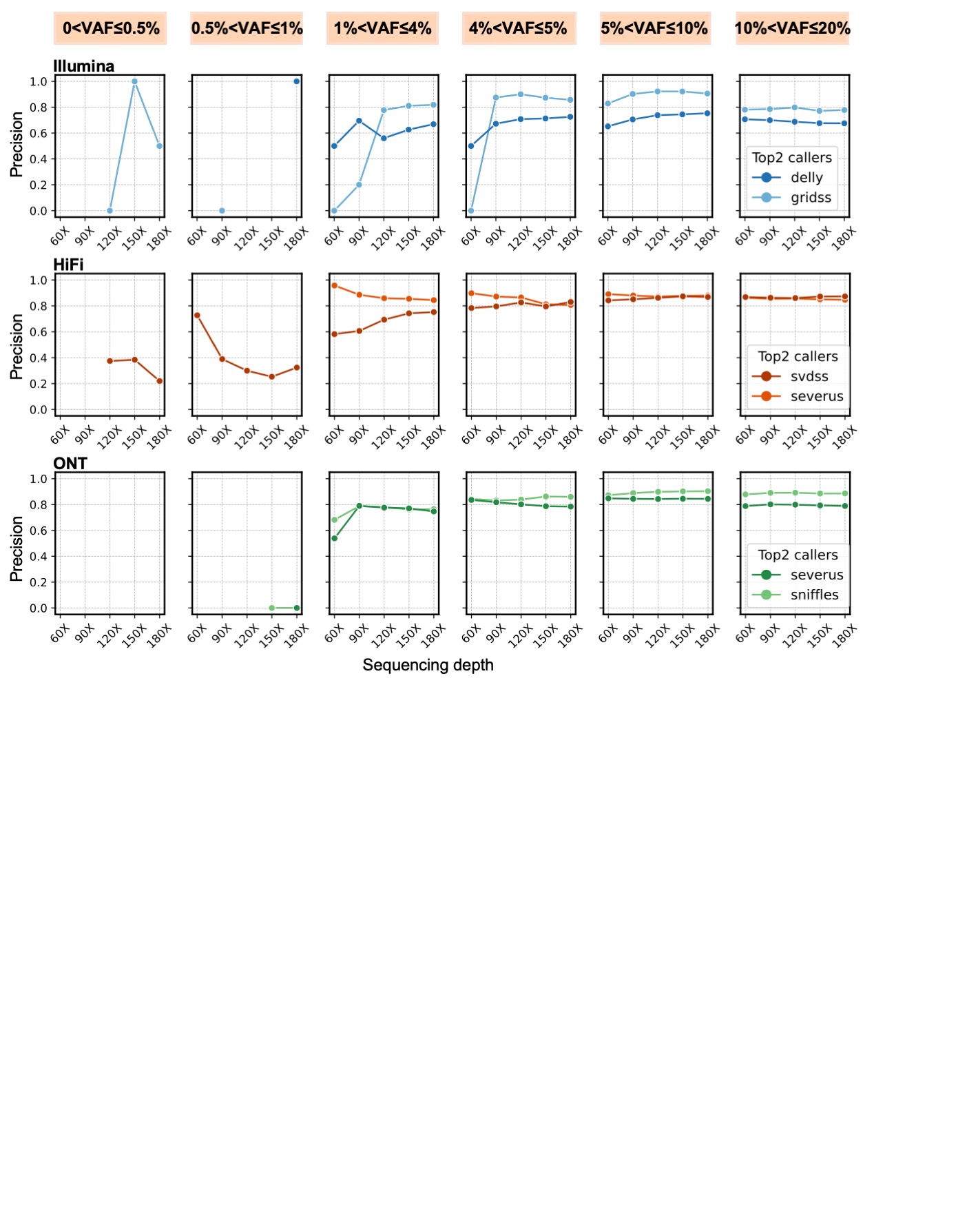

**Supplementary Figure S13 - Precision across VAF bins at fixed sequencing depths.** For each platform (Illumina, HiFi, ONT), the two best callers are shown; each point indicates precision within a VAF bin at the given depth. True and false positives are binned by each caller’s reported VAF. Missing points denote bins with no calls from the caller at that depth (precision is not applicable).

### Section 5

#### StratP Test Stratification Analysis

A selection of 8 SV callers were run on sequencing experiments from 5 GCCs using 3 sequencing technologies and producing 45 VCFs in total. Each of these VCFs were benchmarked against the SMaHT MIMS SV benchmark. The benchmark has 5 main variant features on which variant stratification can be performed: SV Type (N=2); SV Length Bins (N=10); Inside/Outside TR (N=2); Neighboring/Isolated (N=2); VAF Bin (N=4). Combined, these features produce 1,485 possible combinations of variant features. Manually analyzing each feature and its interaction with other features quickly becomes burdensome. Further complicating matters are feature interactions and imbalances where, for example, certain SV types co-occur more frequently within tandem repeats, potentially biasing simple stratification analyses. The StratP test enables statistically rigorous prioritization of features by testing the null hypothesis that a result’s accuracy is independent of a given feature’s value (a.k.a. stratification). A left-tailed StratP test with significance threshold set to 0.01 was performed on the benchmark SVs from each VCF’s truvari result.

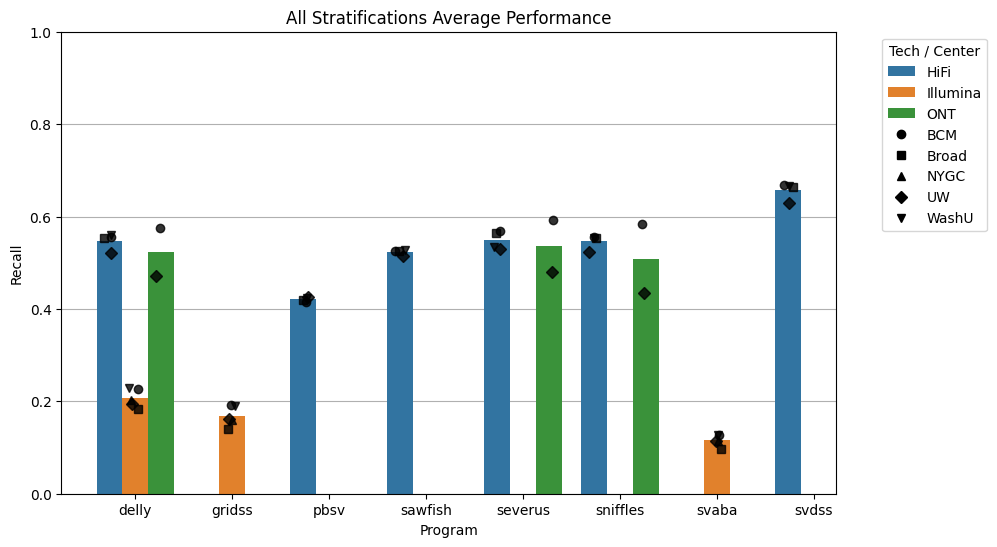

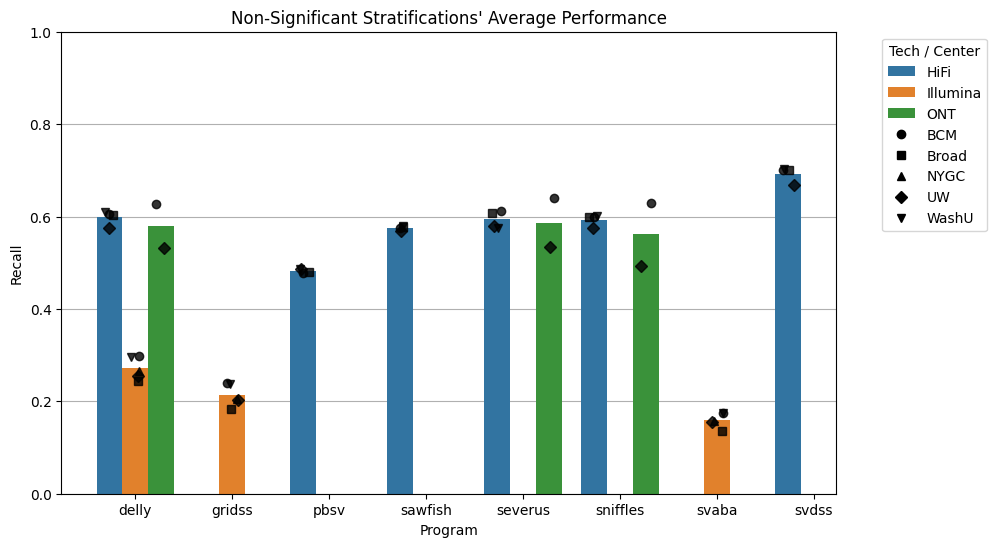

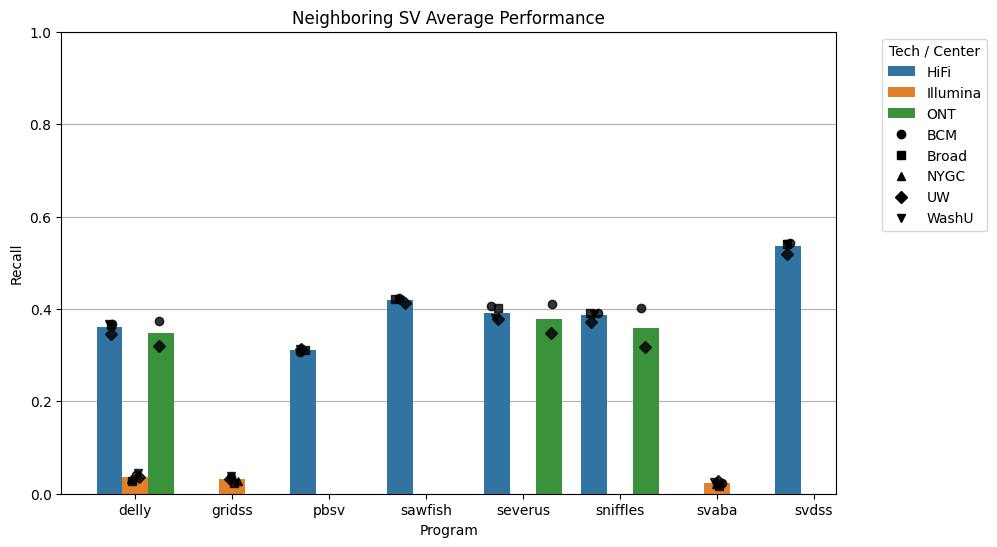

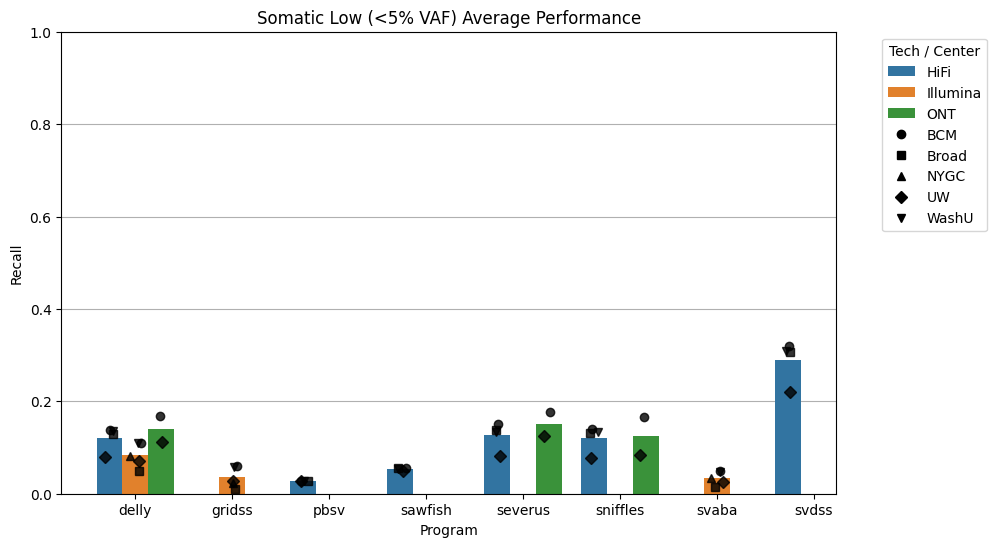

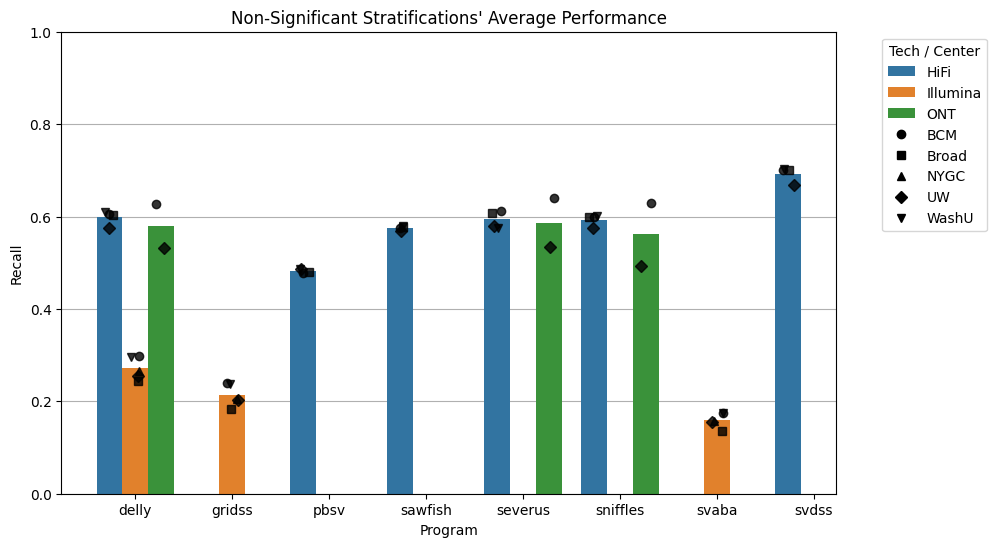

a)

c)

b)

d)

***Supplementary Figure S14 - Average performance across stratifications.*** *a) Mean recall across all 320 stratifications. b) Mean recall across stratifications never identified as having a significant impact on a caller’s results (N=313). c) Mean recall of SVs with Somatic Low <5% VAF per-result. d) Mean recall of SVs with Neighboring SVs within 1kbp per-result.*

Across all stratifications, the average recall was 0.412 (**Supplementary Figure S13a**). Note that this is an averaging of each stratification’s recall (**Supplementary Table 4**), which has redundant counting of variants since they each fall into multiple stratifications (e.g. an SV can be both a deletion and in the [100,200) size bin). Long-read SV callers averaged 0.537 recall whereas short-read callers averaged 0.163 recall. After running the StratP test, 7 stratifications were identified as having a significant impact on at least one result’s recall and were therefore deemed difficult (**Table 1**). The average recall of stratifications which were never identified as impacting a caller’s result was 0.462 (**Supplementary Figure S13b**). The average recall across difficult stratifications was 0.319. Together, the top two difficult stratifications by result total accounted for 59% (20,167 of 34,140) of the benchmark’s SVs.

The stratification which most frequently impacted recall was Somatic Low VAF <5% (**Supplementary Figure S13c**). Only svdss was able to achieve recall above 0.20 on Somatic Low VAF <5% alleles, and did so on all 4 HiFi replicates. The 6 results which did not have a significant dependence on the Somatic Low VAF <5% stratification were all from the Illumina based callers delly (average recall 0.093), and svaba (average recall 0.048). The delly results utilized high coverage sequencing experiments from GCCs WashU (500x coverage), BCM (457x coverage), NYGC (264x coverage), while svaba utilized the same BCM and WashU Illumina sequencing. The higher coverage (≥457x) sequencing experiments being insignificant to svaba while the lower coverage GCC NYGC and Broad (176x) were significant suggests svaba may be more sensitive to coverage than delly. Simultaneously, the StratP test identified Somatic Low VAF <5% as always being a difficulty regardless of coverage for grids, the third Illumina based caller,

The observations of LowVAF having a significant impact on the recall of svdss, with recall ≥0.20, but not on delly’s recall which is lower at 0.093 on Illumina sequencing, exemplify an important property of the StratP test. This test is performed independently on each result. Therefore, a significant stratification to a result should be interpreted as a variant property with which a caller is having difficulties relative to its overall capabilities, and not as a classification of a caller being a worse performer on a stratification relative to another tool.

The second most difficult stratification was Neighboring SVs (≤1 kb from another SV), with 35 of 45 results showing reduced recall (**Supplementary Figure S13d**). The 10 unaffected results were from sniffles (N = 1), svaba (N = 2), sawfish (N = 3), and pbsv (N = 4). Notably, every pbsv result avoided this difficulty, suggesting its algorithm can adequately handle nearby SVs. In contrast, all other callers had at least one result negatively impacted by Neighboring SVs. For these tools’ results where Neighboring SVs were significant, the average recall across callers was 0.267 compared to 0.286 where they were not, indicating that even when not strictly speaking a StratP-identified difficulty, recall remained low and p-values just above significance. This illustrates that the StratP p-value can be used for binary difficult/not-difficult classification, and also to prioritize non-significant stratifications.

For the remaining difficult stratifications, only Illumina based callers had difficulties with Insertions, SVs within TRs, and Small (<200bp) SVs. Separately, 7 long-read results had difficulties with Somatic SVs with VAF between 5%-30%. The first three results were from sniffles, severus and delly, which had difficulties with Somatic 5%-30% VAF SVs on the 65x ONT sequencing experiment from GCC UW (average recall 0.345). However, when these same programs analyzed the 116x ONT sequencing experiment from GCC BCM, their recall no longer had difficulty with this stratification (average recall 0.739). This alludes to the role of coverage for these callers’ ability to detect somatic SVs. Finally, the germline SV caller pbsv had decreased recall of 0.263 on Somatic VAF 5%-30% on all 4 GCC’s HiFi sequencing experiments, significantly lower than its recall of Germline Heterozygous (0.846) and Germline Homozygous (0.931) stratifications.

### Section 6

#### Assessment in other technologies

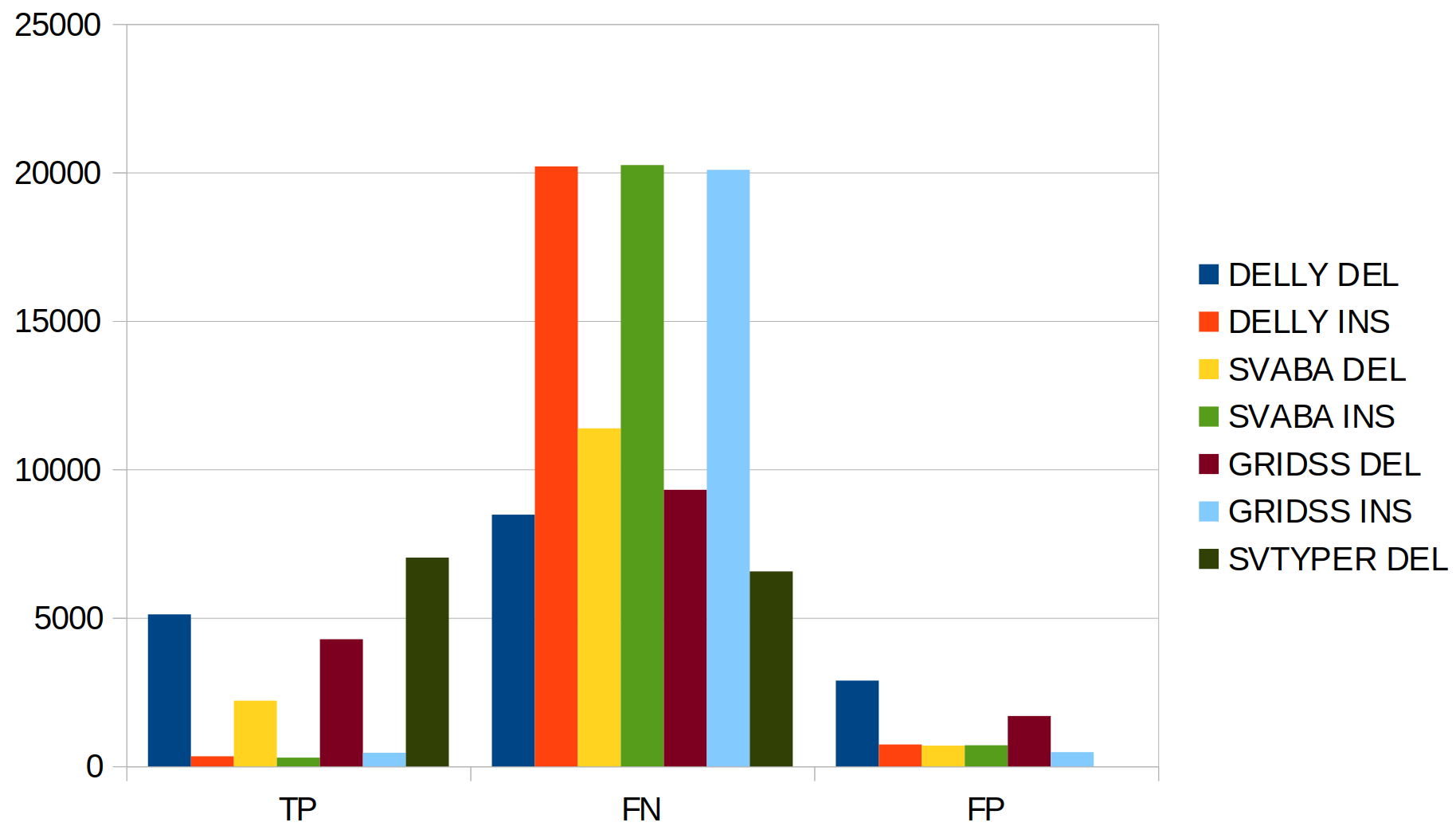

***Supplementary Figure S15*** *- Performance of three short-read SV callers over 232x Element Bioscience Aviti sequencing experiment. Additionally, all DEL were genotyped using SVTyper. Delly shows the highest recall and lowest precision, while SvABA shows the lowest recall and highest precision. Overall, all tools show similar performance in INS. Note: genotyping do not produce FP*

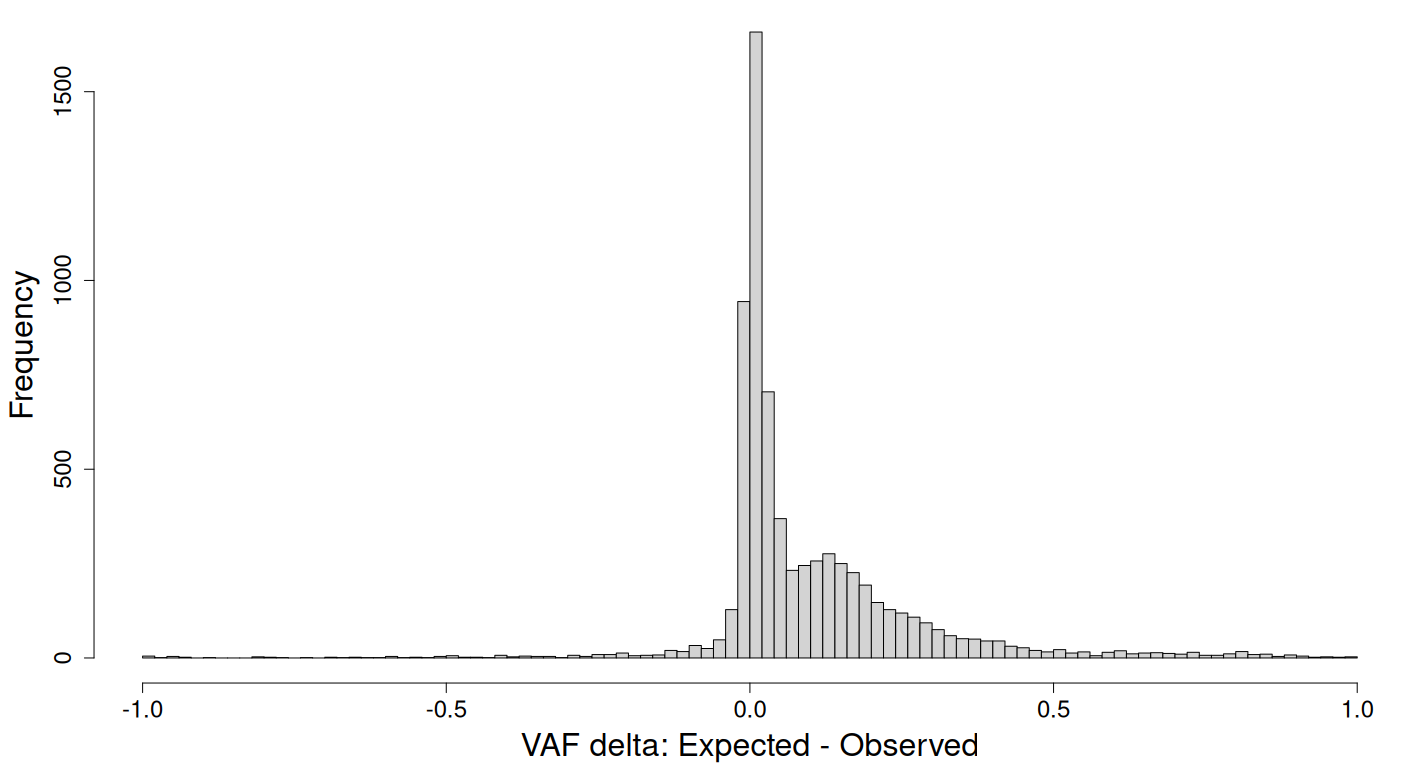

***Supplementary Figure S16*** *- VAF delta of the genotyped DEL. We compared the expected VAF based on the GT and proportion of each sample in the mix to the computed VAF from the genotyper tool. We observed an excess of SVs genotyped with a lower VAF when compared to the benchmark.*

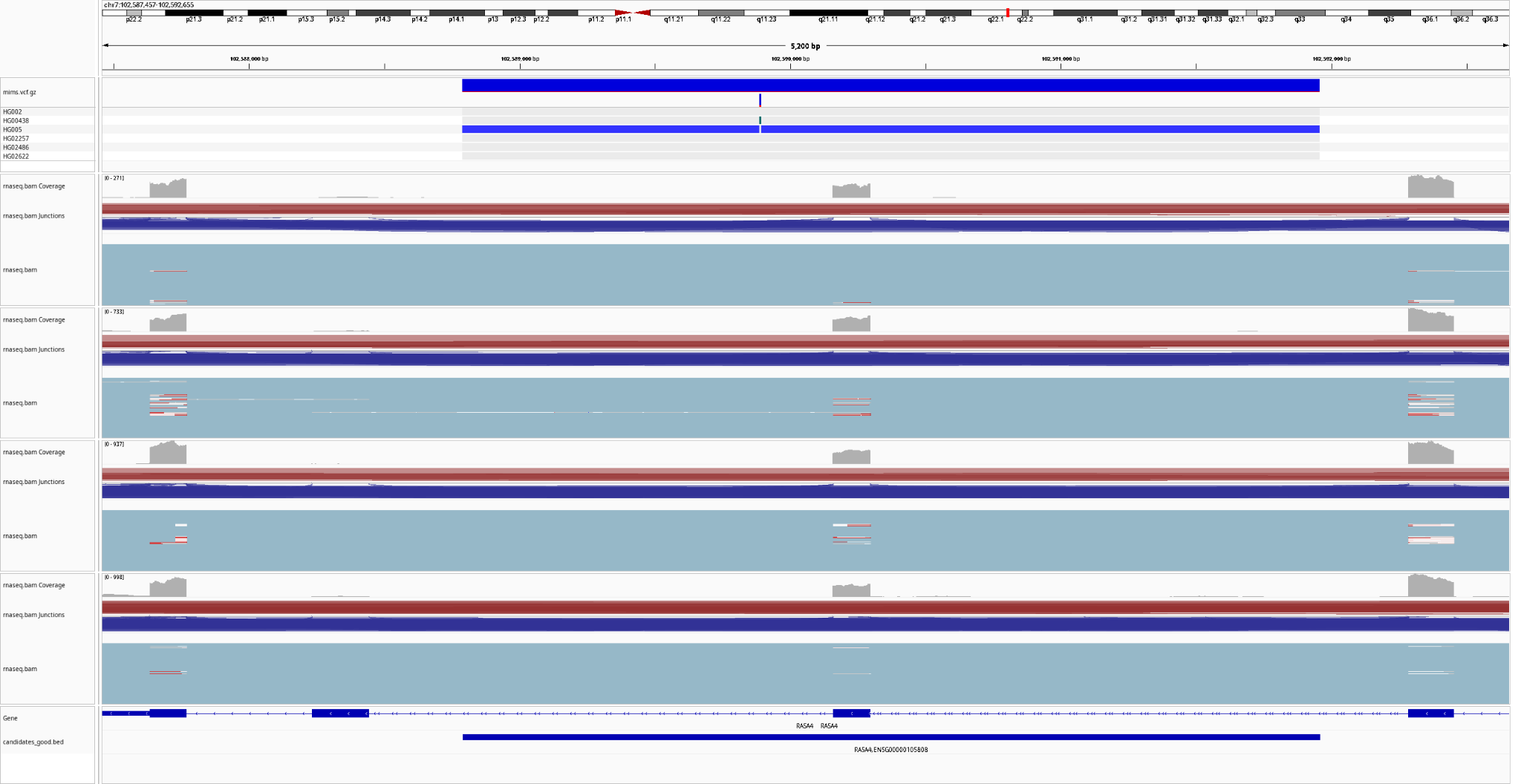

***Supplementary Figure S17*** *- Candidate gene-SV (DEL) with an expected VAF of 41.75% affecting one exon of the RASA4 gene*

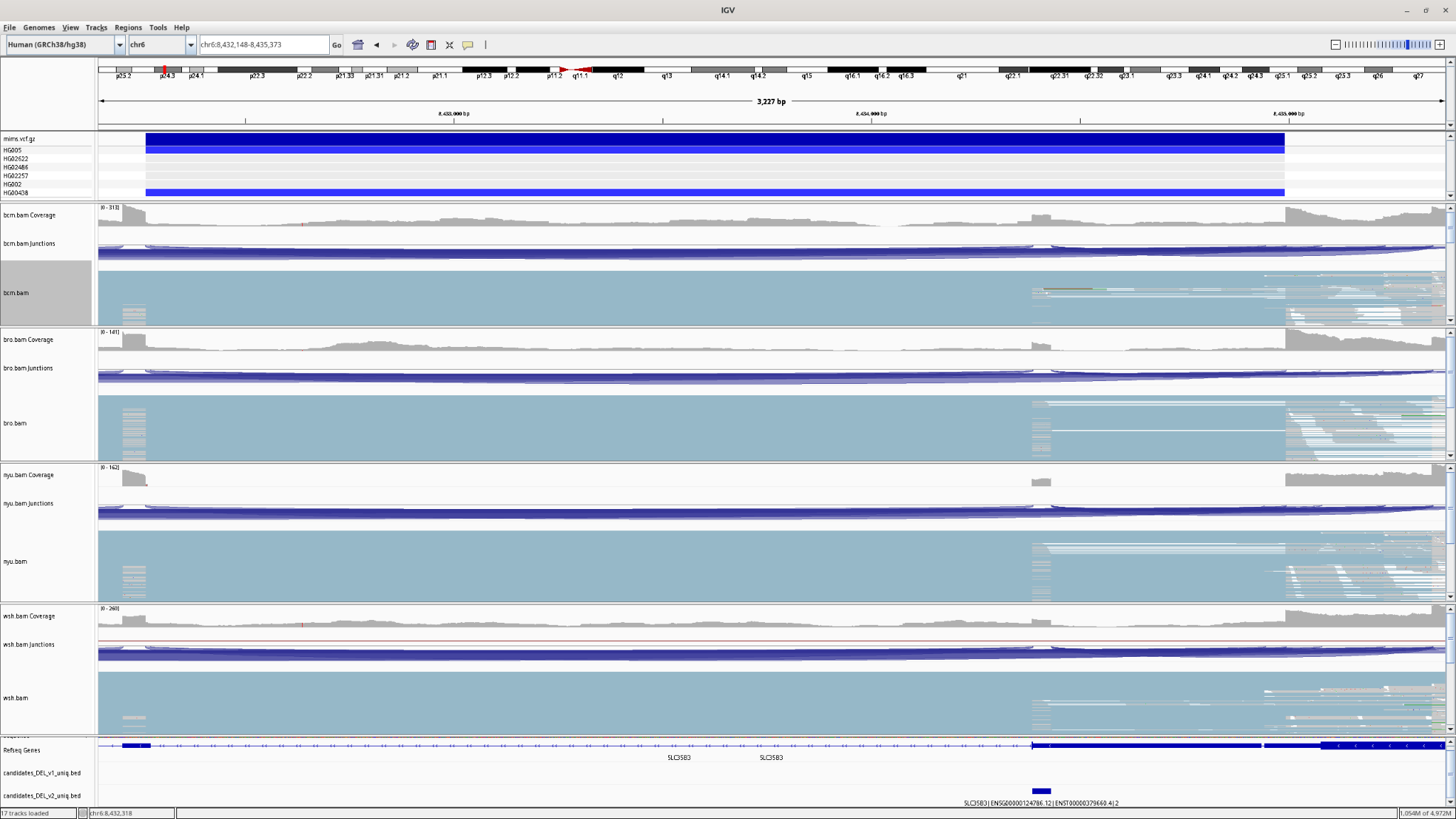

***Supplementary Figure S18*** *- Candidate gene-SV (DEL) with an expected VAF of 42% overlapping with exons of the SLC35B3 gene*

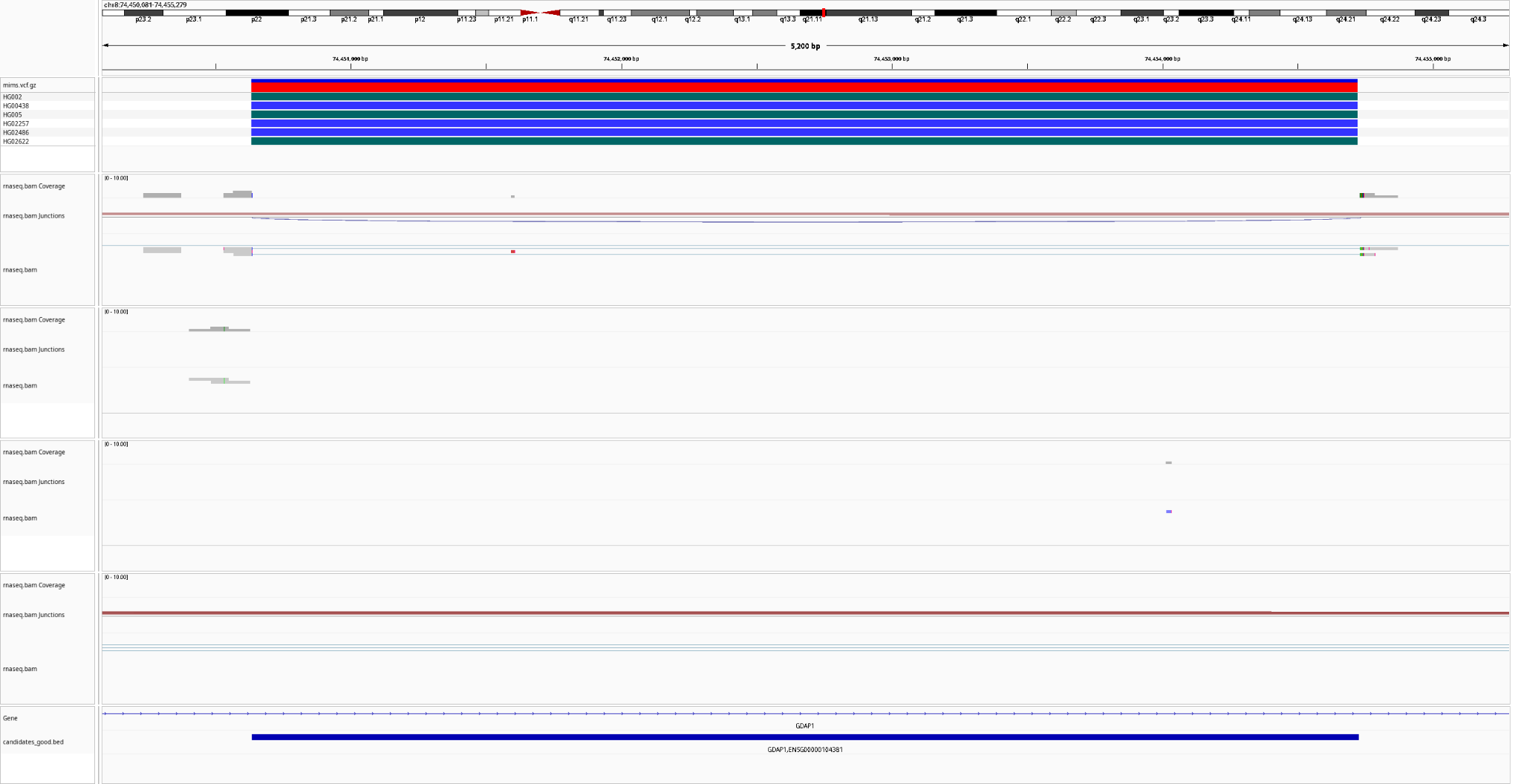

***Supplementary Figure S19*** *- Candidate intron-SV (DEL) with an expected VAF of 93%. It can be seen that the SV slit two reads*

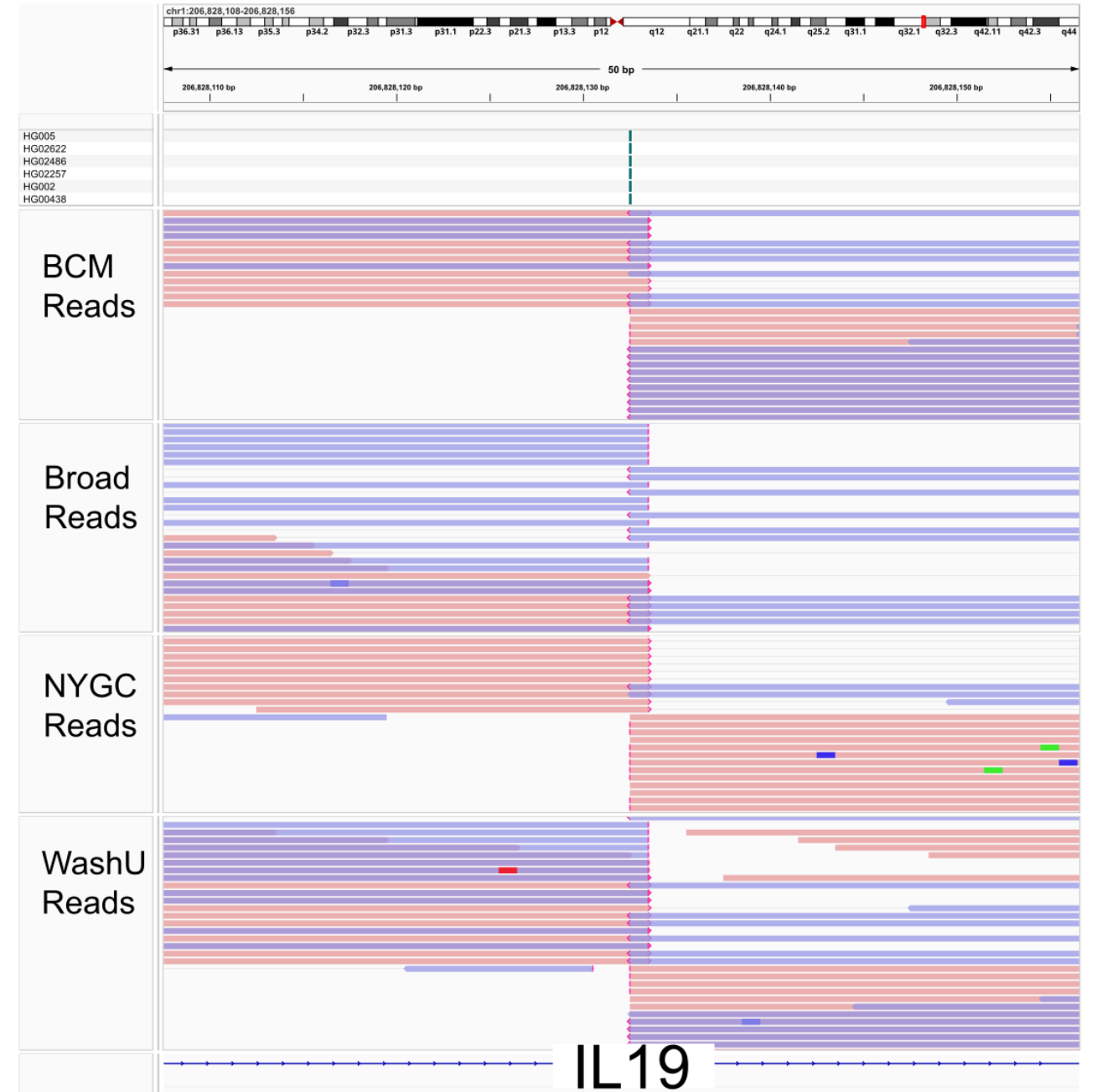

***Supplementary Figure S20*** *IGV screenshot of a germline INS (expected VAF 100%) that overlaps with the IL19 gene. Clear signal from most of the reads which show clipped reads at the exact position of the INS coordinates.*

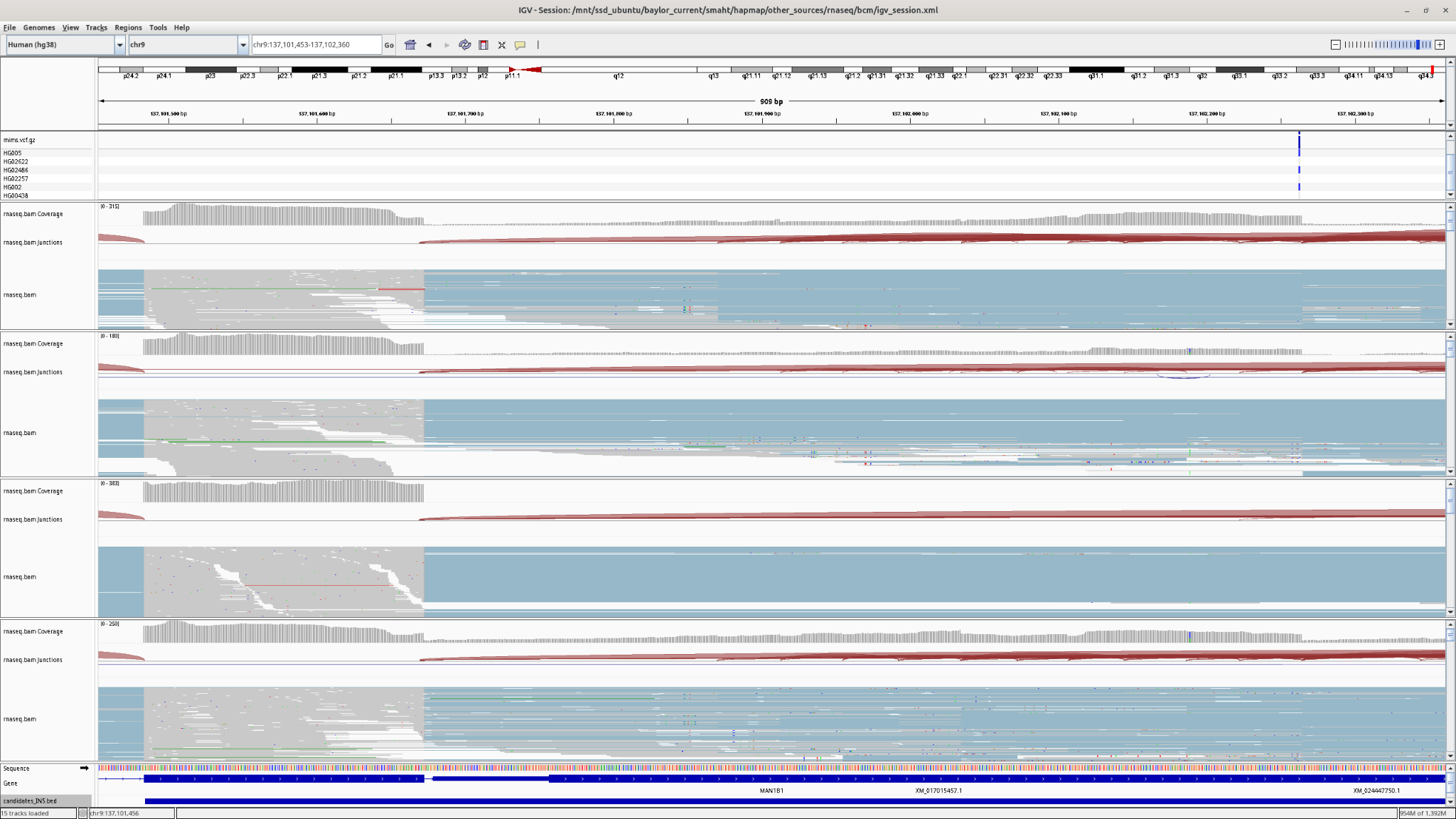

***Supplementary Figure S21*** *- Candidate gene-SV (INS) with an expected VAF of 43.75% .*

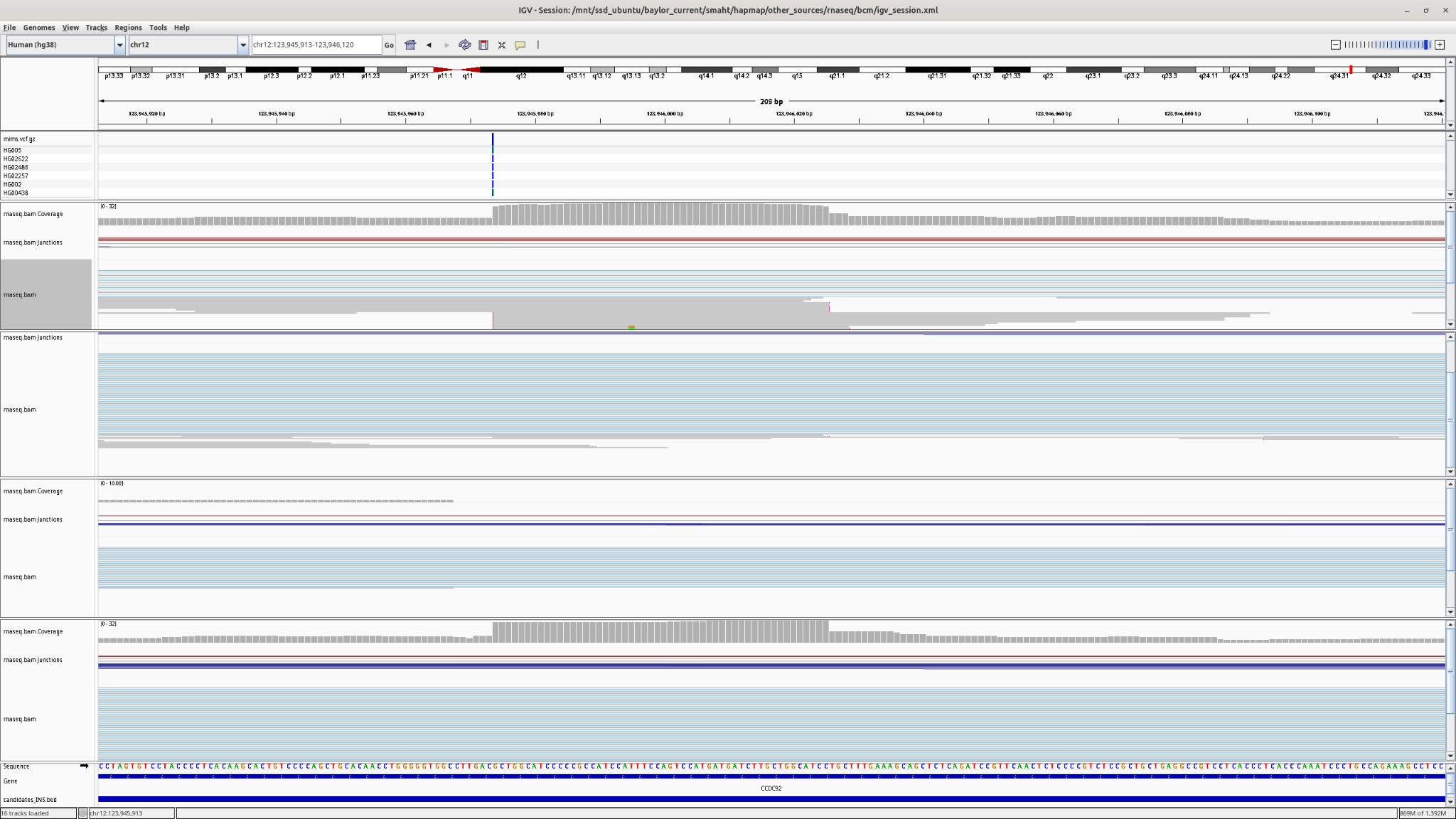

***Supplementary Figure S22*** *-* *Candidate gene-SV (INS) with an expected VAF of 92%*

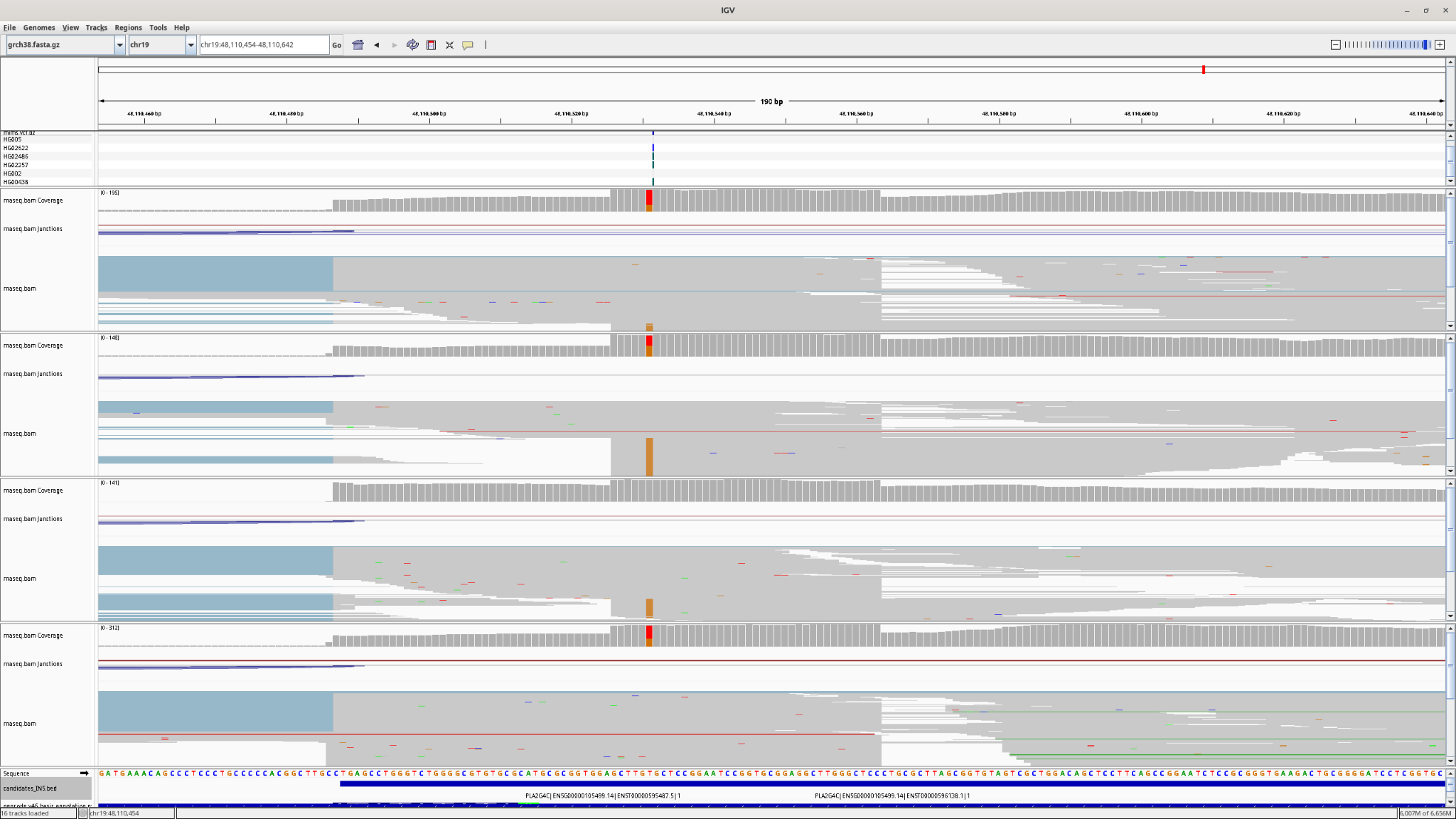

***Supplementary Figure S23*** *- Candidate gene-SV (INS) with an expected VAF of 9.5%*
